# SepF is the FtsZ-anchor in Archaea: implications for cell division in the Last Universal Common Ancestor

**DOI:** 10.1101/2020.10.06.328377

**Authors:** Nika Pende, Adrià Sogues, Daniela Megrian, Hayk Palabikyan, Anna Sartori-Rupp, Martín Graña, Simon K.-M. R. Rittmann, Anne Marie Wehenkel, Pedro M. Alzari, Simonetta Gribaldo

**Affiliations:** Evolutionary Biology of the Microbial Cell Unit, Department of Microbiology, Institut Pasteur, Paris, France; Structural Microbiology Unit, Institut Pasteur, CNRS UMR 3528, Université de Paris, F-75015 Paris, France; École Doctorale Complexité du vivant, Sorbonne University, Paris, France; Archaea Physiology Biotechnology Group, Department of Functional and Evolutionary Ecology, University of Vienna, Wien, Austria; Ultrastructural BioImaging Unit, Institut Pasteur, Paris, France; Bioinformatics Unit, Institut Pasteur de Montevideo, Montevideo 11400, Uruguay

## Abstract

The Archaea present profound differences compared to Bacteria in fundamental molecular and cellular processes. While most Archaea divide by binary fission using an FtsZ-based system similar to Bacteria, they lack the majority of the components forming the complex bacterial divisome. Moreover, how FtsZ precisely functions and interacts with other proteins to assemble the archaeal division machinery remains largely unknown. Notably, among the multiple bacterial factors that tether FtsZ to the membrane during cell constriction, Archaea only possess SepF-like homologues, but their function has not been demonstrated. Here, we combine structural, cellular, and evolutionary approaches to demonstrate that SepF is the FtsZ anchor in the human-associated archaeon *Methanobrevibacter smithii*. 3D super-resolution microscopy of immunolabeled cells shows that *M. smithii* SepF co-localizes with FtsZ at the division plane. We also show that *M. smithii* SepF binds both to membranes and FtsZ, inducing filament bundling. High-resolution crystal structures of archaeal SepF alone and in complex with FtsZ_CTD_ reveal that SepF forms a dimer with a specific homodimerization interface. This drives a strikingly different binding mode from what is observed in Bacteria. Finally, analysis of the distribution and phylogeny of SepF and FtsZ indicates that these proteins date back to the Last Universal Common Ancestor (LUCA) and that Archaea may have retained features of an ancestral minimal cell division system, while Bacteria likely diverged to accommodate the emergence of the complex machinery required to coordinate cytokinesis with the rigid peptidoglycan cell wall and the appearance of additional FtsZ tethers. Our results contribute key insights into the largely understudied mechanisms of archaeal cell division, and pave the way for a better understanding of the processes underlying the divide between the two prokaryotic domains.

## Introduction

Traditionally viewed as extremophiles, the Archaea are now fully recognized as ubiquitous prokaryotes of great ecological and evolutionary importance ^1–3^. Additionally, the human “archaeome” – still in its infancy – is receiving growing attention, as an imbalance of archaeal methanogens has been linked to various pathologies such as Inflammatory Bowel Disease, multiple sclerosis, anorexia and colorectal cancer ^4^. Despite this potential clinical relevance, knowledge on the process of cytokinesis and the involved actors is still very partial in Archaea with respect to Bacteria. While some Archaea (Crenarchaeota) divide by using homologues of the eukaryotic ESCRT system ^5, 6^, most archaeal genomes possess one or two homologues of FtsZ (FtsZ1 and FtsZ2) ^7^. The role of FtsZ during cytokinesis has been shown in a few model archaeal organisms ^8–11^, such as *Haloferax volcanii* where FtsZ1 localizes at the mid-cell constriction site, providing the first cytological evidence that archaeal FtsZ functions in cell division as in Bacteria ^11^. A recent study in *H. volcanii* showed FtsZ1 and FtsZ2 co-localizing at midcell, with both different assembly times and roles for the two proteins^8^.

Only few homologues of the bacterial division machinery that interact with FtsZ have been identified in Archaea, such as the positive and negative FtsZ regulators SepF and MinD, respectively ^7, 12, 13^. SepF was originally identified as a late component of the *Bacillus subtilis* divisome that interacts with the C-terminal domain of FtsZ (FtsZ_CTD_) ^14, 15^. In *B. subtilis*, SepF is non-essential as it has an overlapping role with FtsA, and its deletion only produces aberrant morphologies on the growing septum ^14, 15^. In contrast, in other bacterial lineages where FtsA is absent, such as most *Actinobacteria* and *Cyanobacteria*, SepF is an essential component of the early divisome, orchestrating Z-ring assembly and participating in membrane remodelling ^16–19^. The crystal structure of a bacterial SepF-FtsZ complex was only recently obtained from the Actinobacterium *Corynebacterium glutamicum* ^19^. It showed that SepF forms a functional dimer that is required for FtsZ binding ^19^. *C. glutamicum* SepF binds to the FtsZ_CTD_ through a conserved pocket and interacts with residues at the α-helical interface of the functional dimer ^19^. An early analysis of the distribution of cell division proteins in Archaea revealed that SepF is present in almost all FtsZ-containing taxa, which led to the suggestion that it could act as the main FtsZ anchor ^7^. The structures of two archaeal SepF-like proteins have been reported ^20^, but their biological function has not been studied. Here we provide experimental evidence for a functional role of archaeal SepF in cytokinesis through interaction with FtsZ by using as model *Methanobrevibacter smithii*, the most abundant species of archaeal methanogens in the human microbiome ^4^. We demonstrate that SepF co-localizes with FtsZ to midcell during cell constriction, and binds to membranes. Structural and biochemical analysis shows that SepF also forms a dimer, which binds the FtsZ_CTD_ via a pocket that is partially conserved with Bacteria, but with a markedly different interaction pattern.

We complement these results by a thorough analysis of the distribution and phylogeny of SepF and FtsZ in Bacteria and Archaea. Our results reveal that SepF and FtsZ were already present in the Last Universal Common Ancestor, and that the Archaea might have retained features of an ancestral minimal cytokinesis machinery, while Bacteria diverged substantially, likely to accommodate the emergence of a rigid cell wall and the complex divisome.

## Results

### SepF is widely present in Archaea and co-occurs with FtsZ

We updated the distribution of FtsZ1, FtsZ2 and SepF homologues in the large number of archaeal genomes that have recently become available ^1, 2, 3, 7^ (Materials and Methods, Figure 1, Supplementary Data 1). Most Archaea have one SepF homologue which systematically co-occurs with one or two copies of FtsZ (FtsZ1 and FtsZ2), while it is absent from the genomes that do not have FtsZ homologues – a strong indication of a functional link. Interestingly, while the genes coding for FtsZ and SepF frequently exsist in close proximity in bacterial genomes, this is rarely the case in Archaea, only known to co-occur in a few members of the DPANN superphylum, while they localize with different genes in the other archaea (*ftsZ1* mostly with components of the translation machinery, *ftsZ2* frequently with *parD*, and *sepF* displaying a more variable genomic context) (Supplementary figure 1). Methanopyri are the only Archaea to possess an FtsA-like homologue along with SepF, but its function is unknown. M. smithii contains only one copy of FtsZ1 (*Ms*FtsZ) and one copy of SepF (*Ms*SepF), and therefore being an interesting model with respect to the more complex Haloarchaea which have two FtsZ copies in addition to a large number of FtsZ-like proteins. As no genetic tools are currently available for *M. smithii*, we used an integrative study combining cell imaging, protein biochemistry and structural analysis in order to study the function of SepF.

**Fig. 1.**
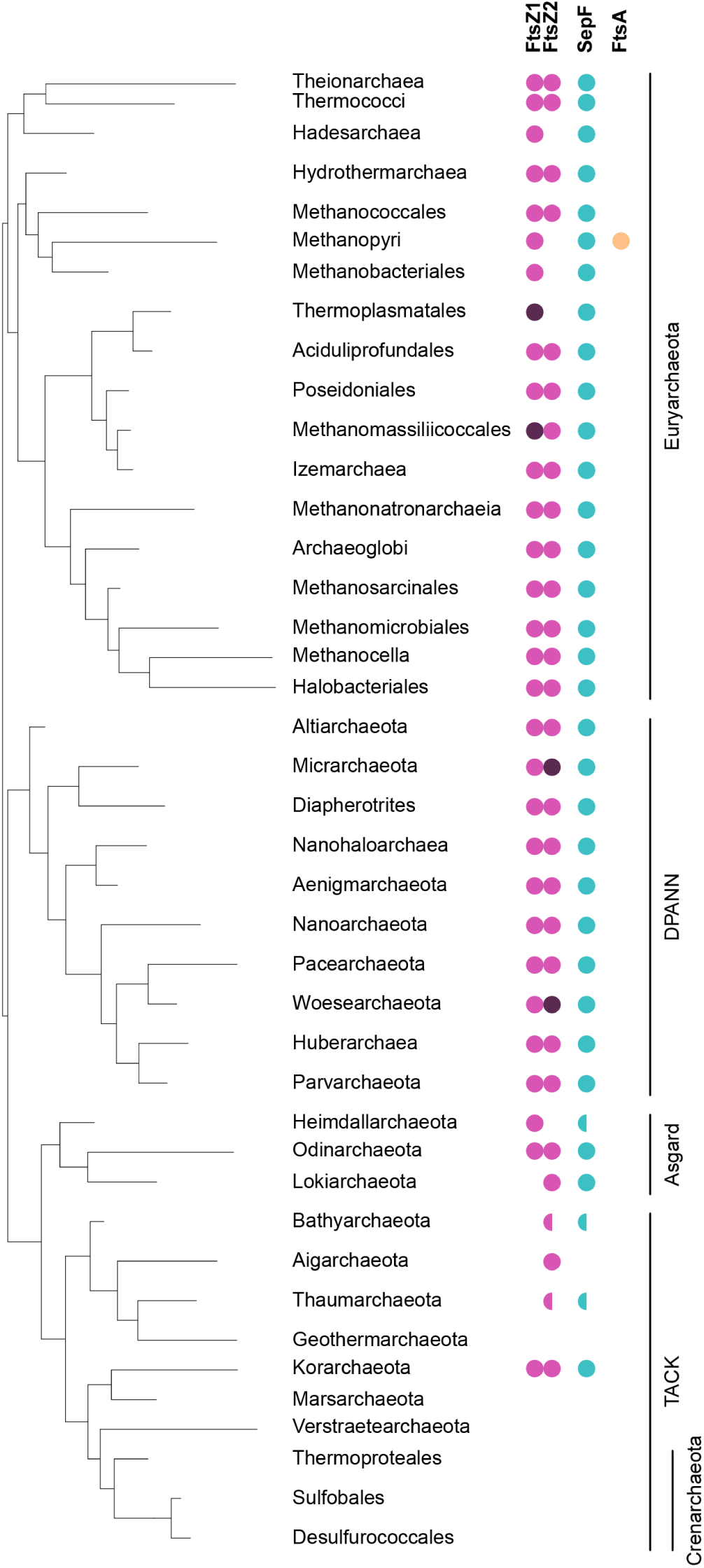
SepF is widely present in Archaea and co-occurs with FtsZ. Distribution of FtsZ1, FtsZ2, SepF and FtsA on a schematic reference phylogeny of the Archaea. FtsZ1 and 2 homologues are present in most archaeal lineages (magenta), and frequently in two or more copies each (dark magenta). Semicircles indicate that the corresponding protein could not be identified in all the taxa of the corresponding clade and may be either due to true absences or partial genomes. The presence of SepF correlates to that of FtsZ in the majority of taxa. An FtsA homologues was only identified in Methanopyri (orange). For full data see Supplementary data 1.

### SepF transiently co-localizes with FtsZ during the *M. smithii* cell cycle

To investigate the function of *Ms*SepF and its potential interaction with *Ms*FtsZ, we studied the localization pattern of these two proteins during the *M. smithii* cell cycle by immunolabelling, using specific anti-*Ms*FtsZ and anti-*Ms*SepF antibodies (Materials and Methods, Figure 2). Since *M. smithii* possesses a cell wall composed of pseudopeptidoglycan (pPG) ^21, 22^ that is resistant to most lysozymes and proteases ^21^, we developed a new protocol for immunolabelling where cells are pretreated with the phage endoisopeptidase PeiW ^23, 24^ which specifically permeabilizes the pPG cell wall (Materials and Methods). Using Western Blot analysis, we quantified FtsZ and SepF levels and showed that the FtsZ/SepF ratio is about six during exponential growth (Supplementary figure 2). To study the intracellular localization of *Ms*FtsZ and *Ms*SepF, we performed co-immunolabelling with anti-*Ms*SepF and anti-*Ms*FtsZ antibodies and imaged the cells by 3-dimensional (3D) super-resolution microscopy (Materials and Methods, Figure 2a). In the ovococcoidal *M. smithii* cells, *Ms*FtsZ forms a discontinuous ring at mid-cell and *Ms*SepF largely overlaps with it (Figure 2a, left panel, Supplementary video 1). At a later stage of division, a smaller FtsZ ring corresponds with the almost completed cell constriction and two new rings appear in the future daughter cells, defining the new septation planes, with SepF forming discontinuous arcs largely overlapping with FtsZ (Figure 2a right panel, Supplementary video 2). Quantitative analysis with several hundreds of co-immunolabeled *M. smithii* cells (Materials and Methods, Figure 2b and c) shows that in cells that exhibiting very slight or no constriction, *Ms*SepF and *Ms*FtsZ co-localize at the future septation plane (Figure 2b, (i) and (ii); 2c, cell 1; Supplementary figure 3). In cells that start to constrict, *Ms*FtsZ is present at the septation plane and *Ms*SepF is found slightly lateral of the Z-ring (Figure 2b, (ii): 2c, cells 2-3, Supplementary figure 3). At a later stage, two distinct fluorescent *Ms*SepF foci can be seen further away from the single *Ms*FtsZ focus (Figure 2b, (iii); 2c, cells 4 - 5; Supplementary figure 3). While *Ms*FtsZ appears restricted to the septation plane throughout the cell cycle, *Ms*SepF is also present lateral of it, which could indicate that SepF moves to the future division planes of the prospective daughter cells before FtsZ. This result agrees with what has been observed in *H. volcanii*, where the 12 amino acids membrane-binding domain of SepF fused to a GFP is able to localize the GFP at midcell in a ring-like structure independently of FtsZ interaction (Nußbaum P., et al., personal communication). Finally, in strongly constricted cells, FtsZ appears in the prospective daughter cells overlapping with the SepF signal, but it is also still present in the current septation plane, where only a weak SepF signal can be detected (Figure 2b, (iv); 2c, cell 6). From these results, we propose a model for FtsZ and SepF localization during the *M. smithii* cell cycle (Figure 2d): (i) in non-constricting cells, both proteins co-localize at the septation plane; (ii) as constriction starts and proceeds, SepF primes the future septation plane in the prospective daughter cells by progressively moving there before FtsZ; (iii) in cells that have almost completed constriction, FtsZ and SepF co-localize at the prospective septation plane of the daughter cells.

**Fig. 2.**
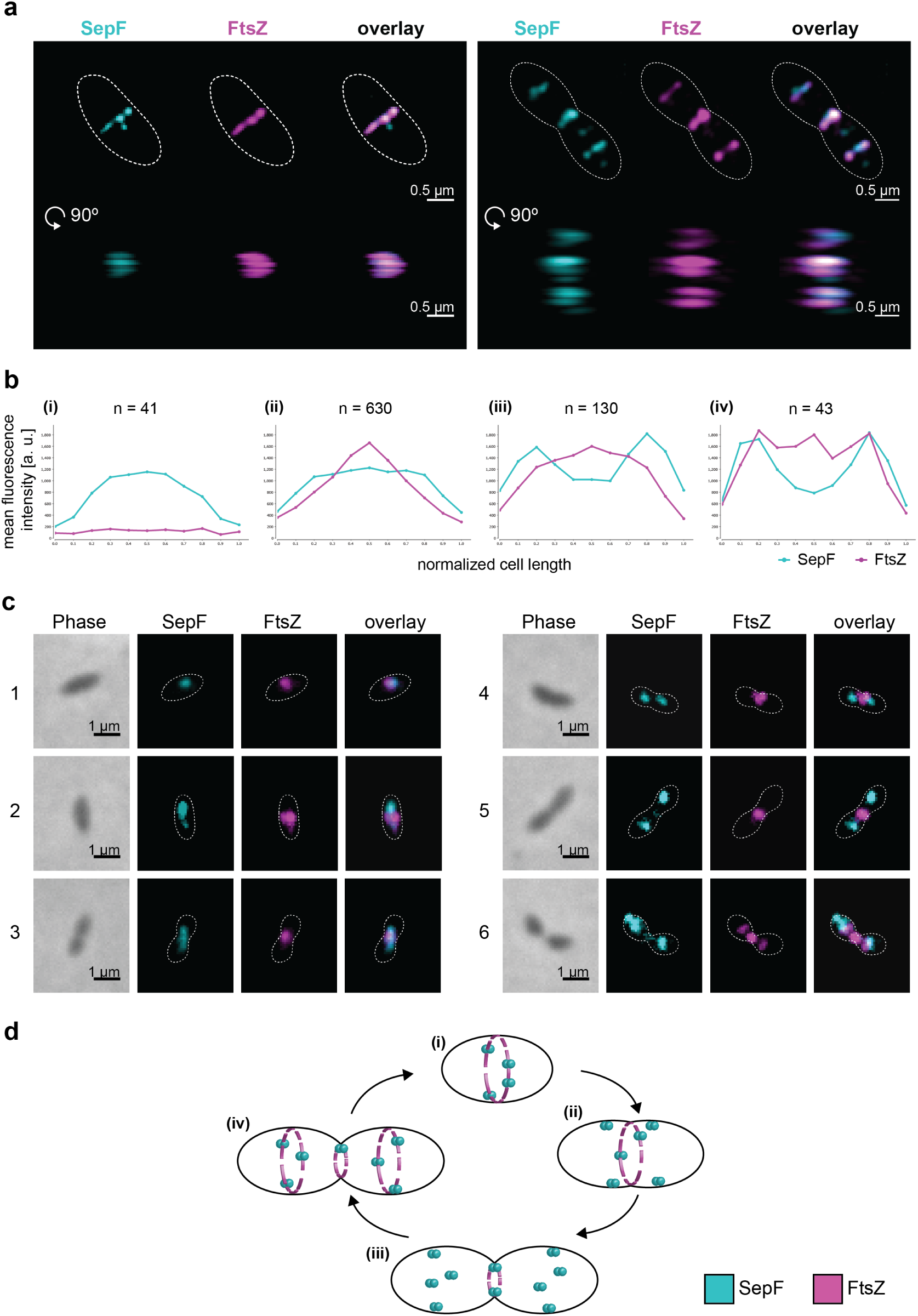
SepF co-localizes with FtsZ during the *M. smithii* cell cycle. *M. smithii* cells were permeabilized with PeiW and immunostained with anti-*Ms*SepF and anti-*Ms*FtsZ antibodies. **a**. 3D Structured Illumination Microscopy (SIM) maximum projections of a non-constricting *M. smithii* cell (left panel) and a constricting cell (right panel) stained with anti-*Ms*SepF (cyan) and anti-*Ms*FtsZ antibodies (magenta). Front views of single channels and the overlay are depicted (above) as well as the side view shifted by 90° (below). White dotted lines represent the cell outlines and scale bars are 0.5 µm. **b.** Mean fluorescence intensity plots of cells grouped into four classes according to the detected FtsZ fluorescent maxima (maxima detected 0-3) with the corresponding SepF (cyan) and FtsZ (magenta) mean fluorescence intensity [a. u.] of each group plotted against the normalized cell length [0^−1^]. **c.** Phase contrast (Phase) and corresponding epifluorescence images of representative co-labelled *M. smithii* cells (SepF in cyan, FtsZ in magenta and an overlay of both). Cells are arranged from non-constricting to constricting from 1 to 6. White dotted lines represent the cell outlines deduced from the corresponding phase contrast images and the scale bars are 1 µm. **d.** Schematic view of SepF (cyan) and FtsZ (magenta) localization pattern during the life cycle of *M. smithii*. (i) in non-constricting cells, both proteins co-localize at the septation plane. (ii) and (iii) SepF progressively moves to the future division plane prior FtsZ. (iv) as constriction is almost completed, FtsZ and SepF co-localize at the prospective septation plane of the daughter cells. The data shown here are representative for experiments performed in triplicate.

### *Ms*SepF binds to membranes and to the FtsZ_CTD_ inducing filament bundling

*Ms*SepF is composed of 149 amino acids and presents the same conserved domain organization as previously described for other bacterial SepF proteins ^19, 20^ consisting of an Nterminal amphipathic helix connected through a flexible linker to a putative C-terminal FtsZ-binding core (Figure 3a, Supplementary figure 4). We show that the membrane-binding domain of *Ms*SepF interacts with small unilamellar vesicles (SUVs) with an affinity of 43±0.2 µmol L^−1^ (Materials and Methods, Supplementary figure 5), a value in the same range as that reported for *C. glutamicum* ^19^. Moreover, thermal shift assays show an important increase of the melting temperature of *Ms*SepF_core_ (amino acids from 54 to 149) (from 70.6 to 82.3 °C) upon addition of *Ms*FtsZ_CTD_ (Supplementary figure 6), indicating a direct interaction between the two proteins. As shown for *C. glutamicum* SepF (*Cg*SepF) ^19^, *Ms*SepF is also able to remodel SUVs, as observed by negative stain EM (Figure 3b, left panel), and this remodelling was reversed upon addition of the FtsZ_CTD_, giving rise to smaller, more regular lipid vesicles (Figure 3b, right panel). Additionally, *Ms*SepF_core_ is capable of inducing bundling of FtsZ protofilaments, giving rise to longer and straighter filaments than FtsZ-GTP alone (Figure 3c and d). Taken together, these results strongly suggest conservation of common functional features between archaeal and bacterial SepF.

**Fig. 3.**
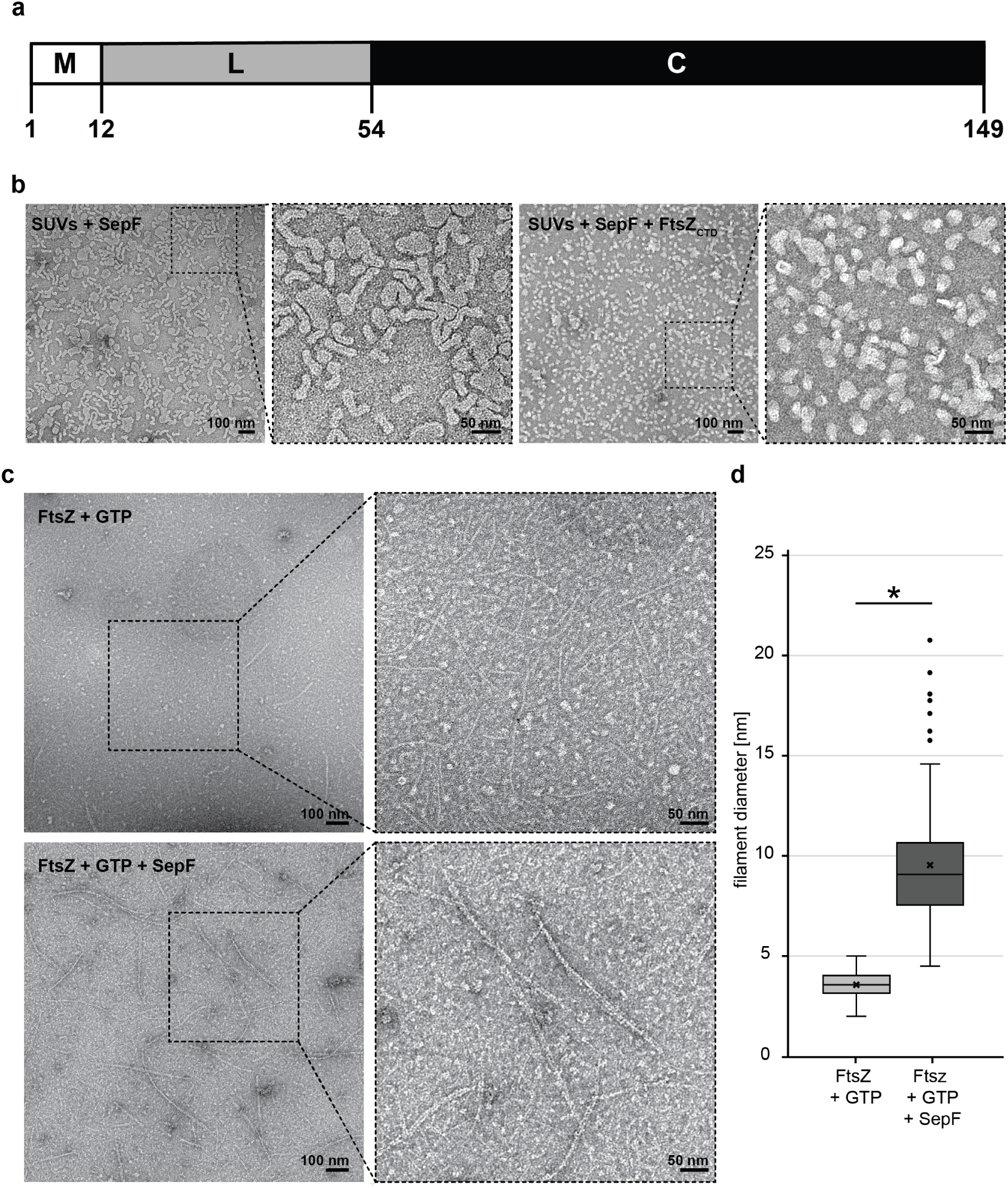
*Ms*SepF binds to membranes and to FtsZCTD inducing filament bundling. **a.** Domain organization of SepF from *M. smithii* with an N-terminal amphipathic helix (M), flexible linker (L) and putative C-terminal FtsZ-binding core (C). **b.** Negative stain electron microscope images of SUVs (100 µmol L^−1^) and *Ms*SepFFL (50 µmol L^−1^) with (left panel) or without FtsZCTD (100 µmol L^−1^) (right panel). **c.** Negative stain electron microscope images of FtsZ (30 µmol L^−1^) and GTP (3 mmol L^−1^) with (upper panel) or without *Ms*SepFcore (20 µmol L^−1^) (lower panel). (**b** and **c**) Panels show original image with an enlarged inlet that is marked by black dotted square next to it. Scale bars are 100 nm (original images) and 50 nm (inlets). **d.** Boxplots showing filament diameter [nm] for FtsZ+GTP and FtsZ+GTP+SepF from 130 filaments each. Box is the inter quartile range, where the lower edge is 25th percentile and the upper edge the 75th percentile. Whiskers show the range between the lowest value and the highest value. Line inside each box indicates the median and (x) indicates the mean. Black circles are outliers. (*) indicates that means of diameter are significantly different.

### High-resolution crystal structures of *Ms*SepF alone and in complex with the FtsZ_CTD_, reveal specific archaeal features

To further characterize the *Ms*SepF_core_, we determined its crystal structure at 1.4 Å resolution (Figure 4a, Supplementary table 1). The protein is a dimer in solution (Supplementary figure 7), with each protomer consisting of a 5-stranded β-sheet flanked by 2 α-helices (Figure 4a). A helical turn (η1) is present between strand β3 and helix α2, which is generally conserved in archaeal sequences but is absent in bacteria (Supplementary figure 8). The overall dimer organization matches that previously described for the structures of archaeal SepF-like proteins from *Archaeoglobus fulgidus* and *Pyrococcus furiosus* ^20^ (Supplementary figure 9), with the dimer interface (1200 Å2) formed by the exposed faces of the two β-sheets, and the last strand (β5) forming part of the opposing beta sheet (Figure 4a). Despite a very similar monomeric structure, the dimer interface in archaeal SepF differs from that observed for the functional dimer in *Cg*SepF, which is formed by a 4 alpha-helical bundle (Figure 4b). Interestingly, both interfaces – formed by the alpha-helices or the beta-sheets – have been observed in crystals of B. subtilis SepF (Figure 4b), where it has been proposed as a putative mechanism of polymerization ^20^.

**Fig. 4.**
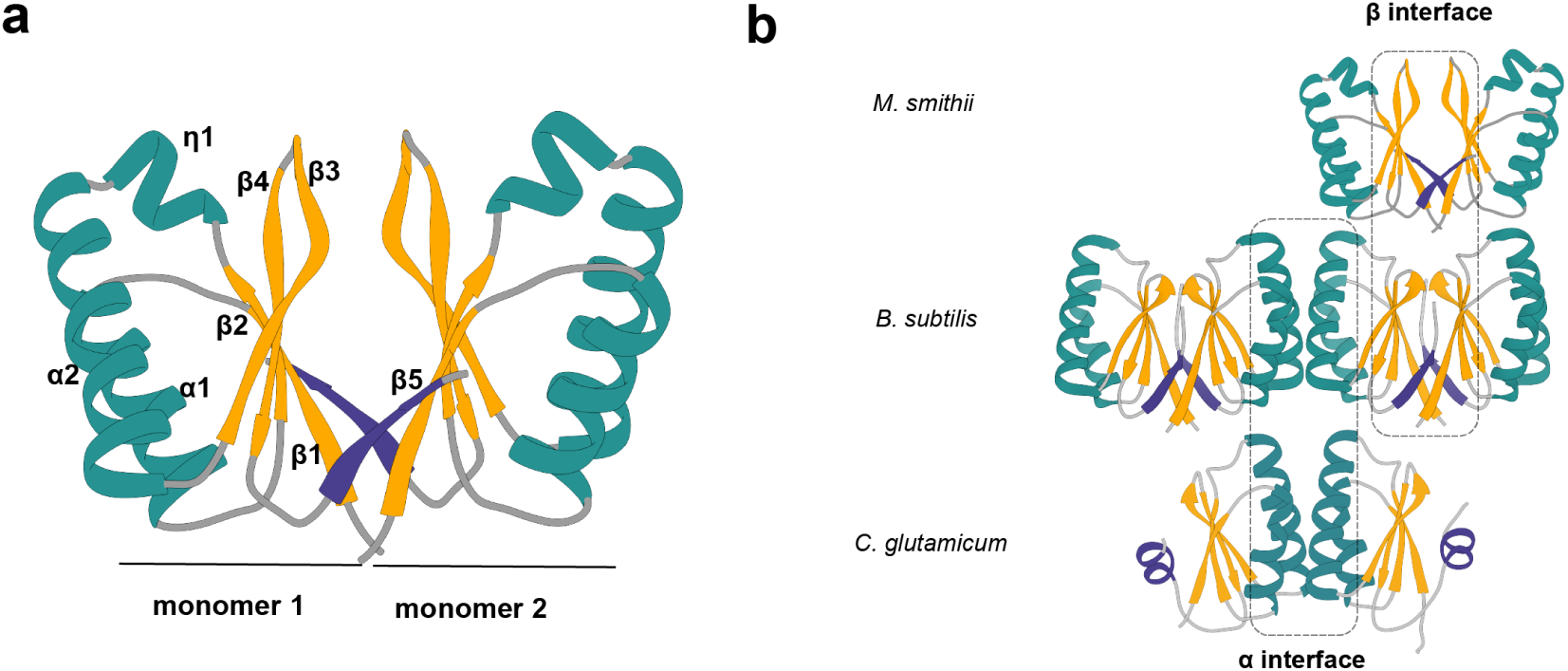
Structural comparison of SepFcore structures. **a.** Crystal structure of the *Ms*SepF dimer composed of two identical monomers. Each protomer consisting in a 5-stranded β-sheet flanked by 2 α-helices and a helical turn (η1). β5 coloured in purple and forms part of the opposing β-sheet. **b.** Comparison of different SepF dimer interfaces in Bacteria and Archaea. The αinterface can only be found in the crystal structures of bacterial SepF such as *C. glutamicum* (PDB 6SCP) and *B. subtilis* (PDB 3ZIH), contrary to the βinterface which is found in all Archaea and *B. subtilis* structures. The C-terminal element of the crystal structures β5 (in both *M. smithii* and *B. subtilis*) and α3 (in *C. glutamicum*) are depicted in purple.

To further study the interaction between SepF and FtsZ, we crystallized *Ms*SepF_core_ in complex with the FtsZ_CTD_ peptide and determined the structure of the complex at 2.7 Å resolution (Supplementary table 1). The bound peptide is well-defined in the electron density for 10 out of 13 residues and adopts an extended conformation within a pronounced groove of the SepF monomer (Figures 5a-b). The association is mediated by both hydrophobic and electrostatic interactions, including three intermolecular salt bridges and 9 hydrogen bonds (Figure 5c). Sequence-based analysis of the FtsZ-binding pocket in archaeal SepF homologues shows general conservation (Figure 5d). Several *Ms*SepF residues that directly interact with FtsZ_CTD_, such as Phe107, Lys115, Arg118, Glu124 and Leu127 are conserved in other archaeal SepF structures (Figure 5c, Supplementary figure 10). The archaeal-specific η1 insertion also contributes to the binding pocket and is in direct contact with the N-terminal glutamine (Gln368) of the FtsZ peptide (Figure 5c). These results indicate that the specific features revealed by our structural data are conserved across the Archaea and are important for FtsZ interaction.

**Fig. 5.**
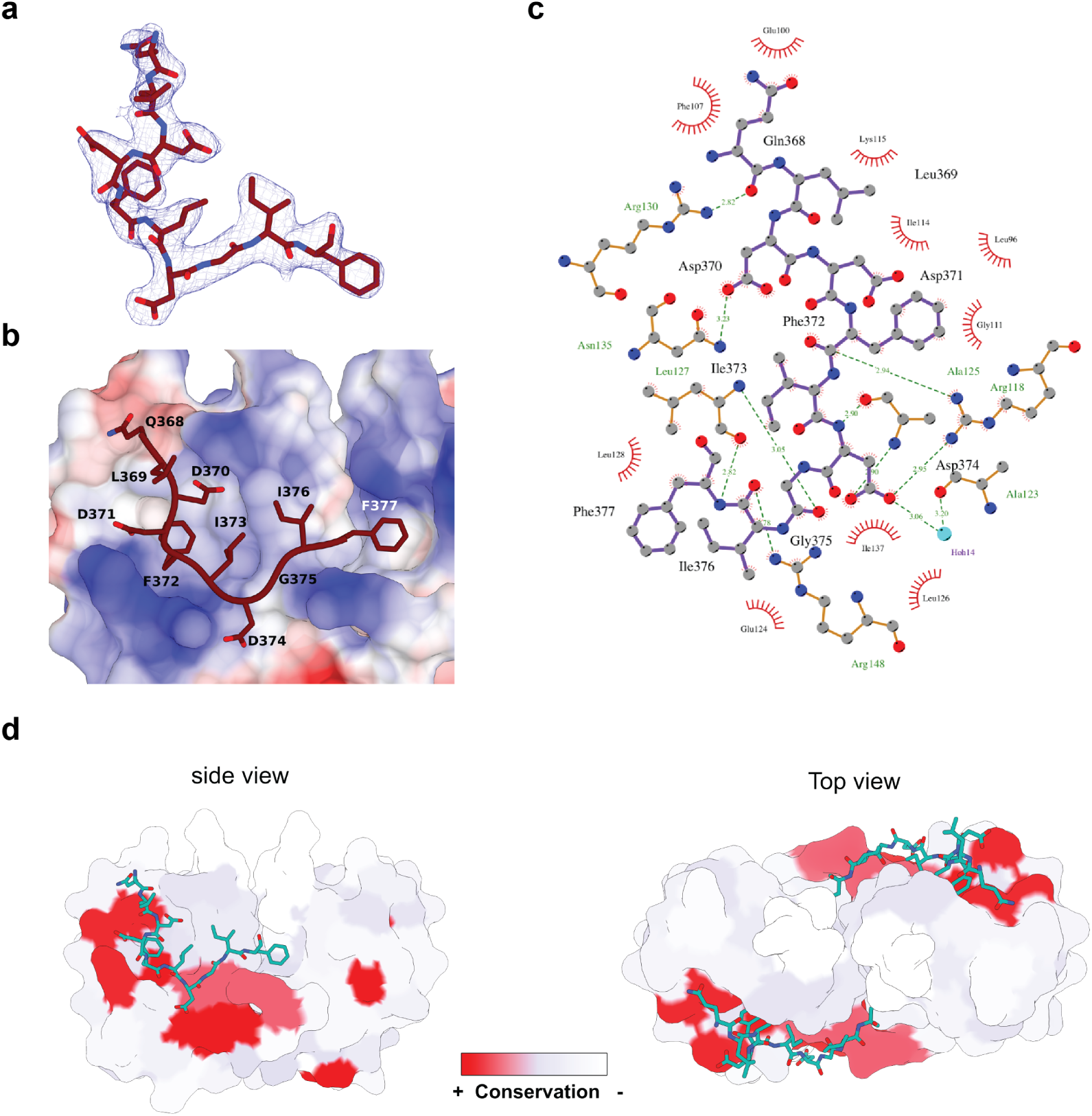
Structural characterization of the *Ms*SepF-FtsZCTD complex. **a.** Final 2Fo-Fc electron density map of FtsZCTD countered at 1.3 σ. **b.** Structure of the FtsZCTD (in stick representation) bound to *Ms*SepFcore binding pocket colorcoded according to surface electrostatic potential. FtsZ residues are labelled. **c**. 2D map of the interactions between the FtsZCTD (purple) and SepF in the biding pocket. Hydrogen bonds are depicted as dotted lines. The plot was made using LIGPLOT 61. **d.** Conservation of the FtsZCTD binding pocket in Archaea. Conserved regions of archaeal SepF sequences were mapped on the *Ms*SepFcore structure using ConSurf-BD 62. Red represents highly conserved residues whereas white indicates poorly conserved. FtsZCTD is shown in turquoise and stick representation.

Despite their evolutionary distance and sequence divergence, the structures of bacterial and archaeal SepF-FtsZ complexes share a conserved mode of interaction for the N-terminal part of the FtsZ_CTD_ (residues LDDFI in *M. smithii* and DLDV in *C. glutamicum*), which binds to a similar groove formed between α2 and β3 in the SepF monomer (Figure 6). However, the binding modes of the C-terminal part of the FtsZ_CTD_ peptide are markedly different between Archaea and Bacteria. In fact, in *C. glutamicum* this region interacts with residues from the two monomers of the SepF functional dimer at the -helical interface (Figure 6 right), while in M*M. smithii* it largely binds to only one monomer of the functional dimer, enlarging the monomer-peptide interface surface (600 Å2) with respect to that observed in *C. glutamicum* (390 Å2) (Figure 6 left). In *M. smithii*, only the terminal residue of FtsZ_CTD_ (Phe377) binds to a surface hydrophobic pocket in the second SepF monomer at the -sheet dimer interface. This residue does not seem conserved in other archaeal SepF homologues except for those phylogenetically close to *M. smithii*. Therefore, the hydrophobic interaction via the FtsZ_CTD_ terminal residue is likely specific to these taxa.

**Fig. 6.**
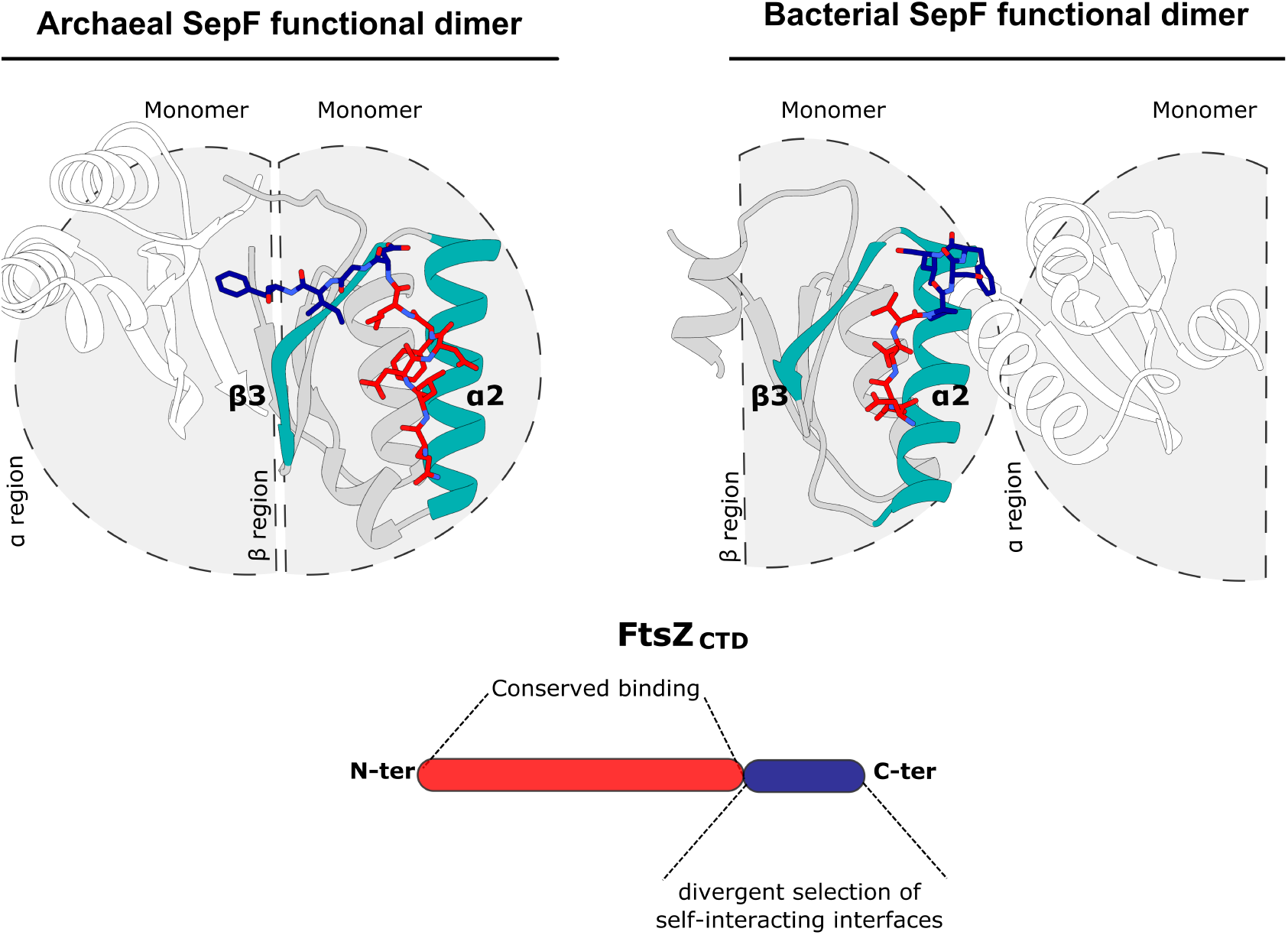
Different SepF functional dimers between Archaea and Bacteria. The self-interacting interface between Archaea and Bacteria that form the functional dimer is different as revealed by the distinct FtsZ-binding modes. The N-terminal part of the FtsZCTD (red) binds to a similar groove formed between α2 and β3 (turquoise) in both SepF monomers, however, the C-terminal part (blue) differs between the two species. In *C. glutamicum* (right) the C-terminal part of the FtsZCTD interacts with residues from the two protomers at the α-helical dimer interface, contrary to *M. smithii* (left) where it interacts largely with only one monomer, and only the last residue (Phe377) interacts with the second monomer at the β-sheet dimer interface. As a consequence, the SepF monomer-peptide interface in *M. smithii* (600 Å2) is higher than in *C. glutamicum* (390 Å2).

### SepF and FtsZ originated prior to the divergence between Archaea and Bacteria

The specificities of archaeal SepF/FtsZ interaction with respect to those observed in Bacteria are intriguing and may lie in an ancient and fundamental divergence in cell division between the two prokaryotic domains. Interestingly, while SepF has a wide distribution in Archaea and is likely the main partner of FtsZ in cell division (Figure 1), its presence in Bacteria is much more scattered, being present mostly in a deep-branching few phyla belonging to the Terrabacteria clade (Actinobacteria, Cyanobacteria, Firmicutes, Fusobacteria, Armatimonadetes), frequently together with FtsA, while it is largely replaced by FtsA in most phyla belonging to the Gracilicutes (including *Escherichia coli*) (Supplementary figure 11 and Supplementary data 2).

The phylogeny of SepF clearly shows a separation of archaeal and bacterial homologues and – although not fully resolved due to the small number of informative positions – an overall topology consistent with vertical inheritance, i.e. monophyly of major phyla and relationships roughly congruent with the reference bacterial phylogeny (Figure 7a). A similar evolutionary history can be inferred from FtsZ, which shows a clear separation among archaeal and bacterial homologues (Supplementary figure 12), and suggests an ancient gene duplication in Archaea giving rise to FtsZ1 and FtsZ2 paralogues, consistent with recent reports ^8^. These results strongly suggest that SepF and FtsZ were likely part of the division machinery in the Last Universal Common Ancestor (LUCA) and co-evolved in the two prokaryotic domains. FtsA phylogeny indicates that it likely arose early on in Bacteria (Supplementary figure 14) and gradually took over SepF function, which was kept in some phyla but otherwise massively lost during bacterial evolution. In contrast, SepF was retained in Archaea, and the FtsZ1 and FtsZ2 paralogues were either kept or differentially lost during archaeal diversification(Figure 7b).

**Fig. 7.**
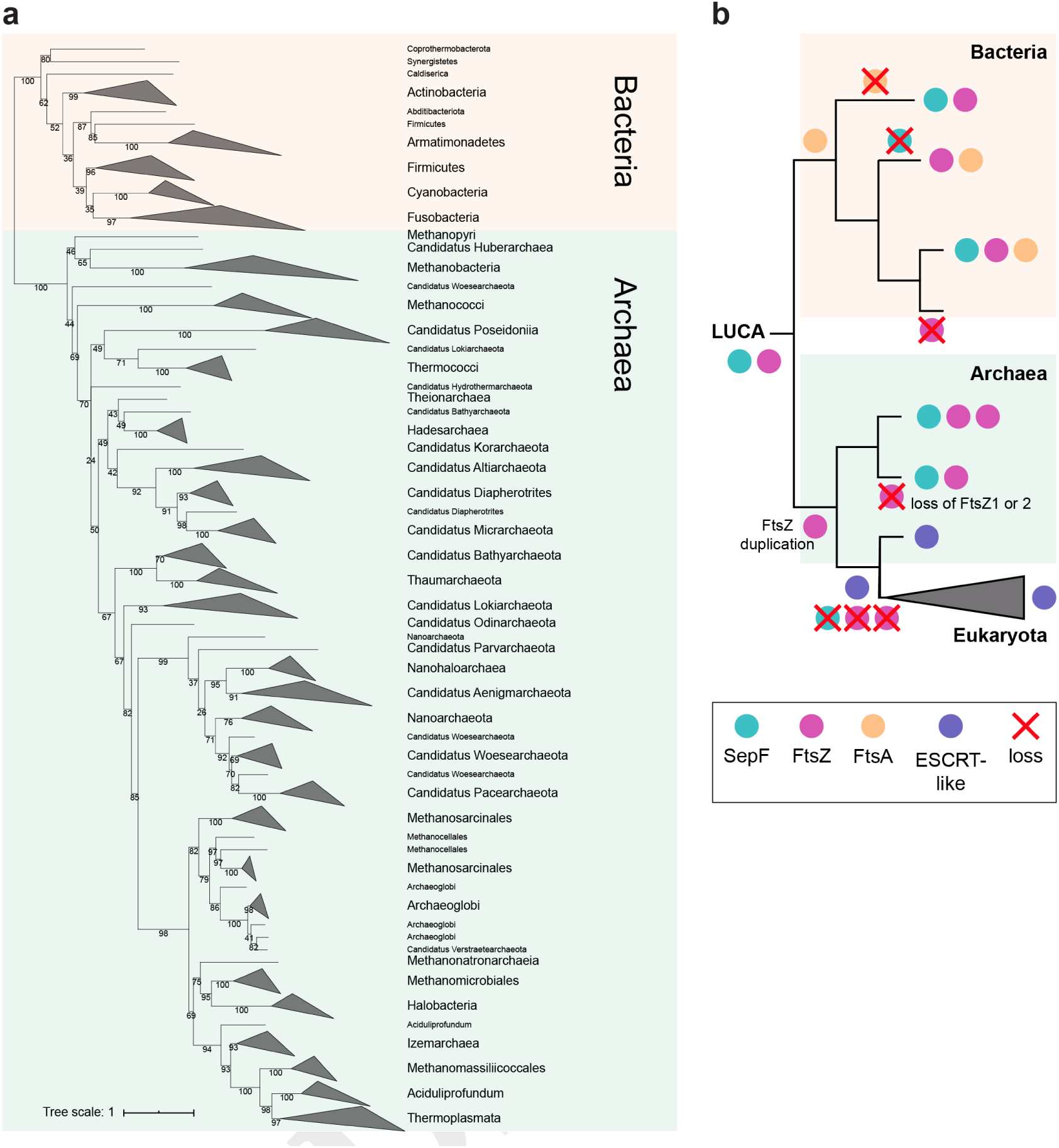
Phylogeny of SepF homologues in Bacteria and Archaea. **a.** Maximum likelihood tree of SepF inferred with IQ-TREE v1.6.7.2 (ModelFinder best-fit model LG+F+R5) ^55, 57^ from an alignment of 147 sequences and 143 amino acid positions. Numbers at nodes represent ultrafast bootstrap supports ^56^. The scale bar represents the average number of substitutions per site. There is a clear separation between Bacteria and Archaea and the tree roughly recapitulates known phylogenetic relationships. This result suggests that SepF was already present in the LUCA and followed a mainly vertical evolution in the two prokaryotic domains. While SepF was lost in many Bacteria, it was largely retained in Archaea. **b.** Schematic scenario for the origin and evolution of FtsZ-membrane tethering in Bacteria and Archaea.

## Discussion

Cell division is one of the most ancient and fundamental processes of life. Yet, Bacteria and Archaea present profound differences in the way they divide, illustrated by the fact that the majority of components of the bacterial divisome seem to be absent in Archaea. In contrast to the well-studied FtsA, the function of SepF has been analysed in a few bacteria so far ^14, 15, 17, 18, 19, 20, 25^, and only one crystal structure in complex with FtsZ is currently available from *C. glutamicum* ^19^.

Here, we report the first experimental evidence that archaeal SepF also acts as the membrane anchor during cytokinesis. By applying our newly developed immunolabelling protocol for archaeal methanogens with a cell wall made of pPG and 3D super resolution microscopy, we show that the only copy of FtsZ in *M. smithii* forms a ring-like structure and colocalizes at mid cell with SepF. A very recent study in *H. volcanii* suggests that FtsZ1 goes first to the division plane followed by SepF that then recruits and anchors FtsZ2 to the septum (Nußbaum P., et al., personal communication). In *M. smithii* SepF localizes to the future division site before FtsZ, thereby priming the future septation plane (Figure 2d). This is in contrast to what was observed in the ovococcoid bacterium *Streptococcus pneumoniae*, where SepF and FtsZ initially colocalize at the septum, but then FtsZ only relocalizes to the new division sites ^26^. However, *S. pneumoniae* also possess the essential alternative Z-ring tether FtsA 26 as well as MapZ, a transmembrane protein that forms a ringlike structure, marks the division site before FtsZ and positions the Z-ring ^27^. It is possible that in *M. smithii* SepF also has a role similar to bacterial MapZ in priming the division site and placing FtsZ, but this will need further investigations. Our results show that SepF-mediated anchoring of FtsZ to the membrane for cytokinesis dates back to billions of years ago, prior to the divergence between Archaea and Bacteria. In agreement with this hypothesis, the crystal structures of the SepF-FtsZ_CTD_ complex reveal an ancestral binding mode that is conserved in the currently available structures from Bacteria ^19^ and Archaea (this work), although the consensus motif of the archaeal FtsZ_CTD_ is shorter than its bacterial counterpart (Supplementary figure 13). However, despite this functional conservation, the crystal structures of archaeal SepF alone and in complex with the FtsZ_CTD_ reveal clearly distinctive features, notably at the level of the dimer interface and FtsZ binding. Such major divergence may be linked to fundamental differences in cell envelope and division in the two prokaryotic domains which are worth discussing.

The overall structure of the SepF monomer is very similar in all available archaeal and bacterial crystal structures (Supplementary figure 9), and in all cases the functional unit is a dimer. However, the functional dimer (as defined by its complex with FtsZ) strikingly differs between *Ms*SepF, where it is formed via a β-βinterface, and *Cg*SepF, where it is formed via a α-αinterface (Figure 4b and Figure 6). Importantly, a bacterial-like α-αinterface is not possible in archaeal SepF because a crucial glycine residue (Gly114 in *Cg*SepF) buried at the center of the bacterial interface is substituted by charged residues with longer side chains (Lys, Glu, Asp) in archaeal sequences (Supplementary figure 10), precluding such interaction to occur. Therefore, the very different ways in which *Ms*SepF and *Cg*SepF bind to FtsZ (by each monomer or in the functional dimer interface, respectively) are driven by their specific self-association modes.

Interestingly, the crystal structure of *B. subtilis* SepF has shown that it can form both bacterial-like α-αand archaeal-like β-βinterfaces ^20^ (Figure 4b). The presence of these two self-interacting interfaces has suggested a simple mechanism for *B. subtilis* SepF, where FtsZ binding occurs through the α-αinterface and polymerization through the β-βinterface ^20^, a mechanism which would therefore be absent in Archaea. However, the only other available bacterial SepF structure (*Cg*SepF) revealed that a C-terminal helix α3 interacts with the β-sheet of SepF and would therefore preclude formation of a β-βinterface ^19^. Interestingly, helix α3 is predicted to be conserved in most bacterial sequences including *B. subtilis*, but not in Archaea (Supplementary figure 8). This observation may suggest that the β-βinterface observed in the crystal structure of *B. subtilis* SepF is not involved in bacterial SepF polymerization and might rather be a vestigial feature of the ancestral archaeal-like SepF. Alternatively, by interacting with other cell division partners, the amphipathic helix α3 may be involved in regulating bacterial SepF polymerization through formation of a β-βinterface, a possibility that will require further study.

In Bacteria, SepF polymerization has been associated with membrane remodelling ^19, 20^. Here, we show that *Ms*SepF is also able to tubulate liposome surfaces, although it remains to be determined whether this requires polymerization. In fact, we (and others, Nußbaum P., et al., personal communication) did not obtain conclusive experimental evidence that *Ms*SepF polymerizes, and the three available archaeal SepF structures provide no hints about a possible mechanism.

The fundamental divergence between SepF/FtsZ interaction in *M. smithii* and *C. glutamicum* correlates well with the profound differences in the cell envelope structure of Archaea with respect to Bacteria, the former being much more variable and generally lacking a peptidoglycan cell wall ^22^. In contrast, the emergence of a rigid cell wall made of peptidoglycan occurred very early in bacterial evolution and likely drove the emergence of the associated complex multi-protein divisome machinery to provide additional mechanical force. Our results suggest that FtsA has also an early origin in bacteria (Supplementary figure 14), and that it may have provided an alternative tether, eventually replacing SepF during bacterial diversification. This phenomenon could be linked with the fact that FtsA is a highly dynamic ATP-dependent protein that might therefore have allowed to couple cell division with the energy status of the cell.

The evolution of cytokinesis in Archaea seems to have been less constrained and more dynamic, with (1) the early presence of two FtsZ paralogues and their subsequent independent losses ^1, 7^, (2) the appearance of a multitude of FtsZ-like protein families in some lineages such as Haloarchaea ^28, 29^, and (3) eventually the emergence of the ESCRT-like system and its takeover of cytokinesis in the archaeal lineages closest to the origin of eukaryotes ^1, 7^ (Figure 7b). The emergence of a cell wall made of pPG in *M. smithii* and all *Methanobacteriales* and the related *Methanopyrales* is more recent than in Bacteria and is very likely an evolutionary convergence. It will be interesting to investigate if it led to a peculiar SepF/FtsZ interaction by obtaining additional structural data in other wall-less archaeal lineages. Indeed, a single amino acid (Phe377) in FtsZ specifically interacts with the second protomer in the SepF dimer (Figure 5). This position is only conserved in *M. smithii* related taxa and perhaps provides extra mechanical strength to drive invagination of their thick pPG cell wall. Finally, our results strongly suggest that a SepF/FtsZ system was already present in the LUCA, unequivocally defining it as a cellular entity. LUCA may not have possessed a complex envelope, and cell division could have been mainly achieved by a minimal system where SepF converted the dynamic energy of FtsZ into mechanical force to achieve membrane constriction ^30, 31^. Several pieces of evidence suggest that the LUCA only contained a rather basic plasma membrane with the absence of a cell wall ^32, 33^, thus, a membrane-interacting protein bound to a dynamic cytoskeleton element could have represented enough to promote cell division. Therefore, a plausible evolutionary scenario could be that SepF and FtsZ represent the minimal ancestral cell division apparatus in the LUCA, possibly together with a few auxiliary proteins. Given the large conservation of the SepF/FtsZ system in Archaea with respect to Bacteria, combined with its peculiar features, it is tempting to speculate that it may have retained ancestral features dating back to the LUCA.

In conclusion, our results pave the way for future studies to understand cell division in different archaeal models, in particular gut-associated methanogens with a cell wall made of pPG. FtsZ/SepF interaction represents only part of the story and many questions remain to be answered, such as how the Z-ring is positioned in Archaea, and which other proteins are part of the archaeal divisome. The answers to these questions will allow to reveal fundamental features of contemporary archaeal cell biology, while at the same time dwelling into the most ancient evolutionary past.

## Material and Methods

### Bacterial and archaeal strains and growth conditions

All bacterial and archaeal strains used in this study are listed in Supplementary table 2. *Escherichia coli* DH5αor Top10 were used for cloning and were grown in Luria-Bertani (LB) broth or agar plates at 37 °C supplemented with 50 µg mL^−1^ kanamycin or 100 µg mL^−1^ ampicillin when required. For protein production, *E. coli* BL21 (DE3) was grown in LB or 2YT broth supplemented with 50 µg mL^−1^ kanamycin or 100 µg mL^−1^ ampicillin at the appropriate temperature for protein expression. *Methanothermobacter wolfeii* DSM 2970 and *Methanobrevibacter smithii* strain DSM 861 were obtained from the Deutsche Sammlung von Mikroorganismen und Zellkulturen (DSMZ; Braunschweig, Germany). *M. wolfeii* and *M. smithii* were grown approximately two to three weeks in 100 mL serum bottles (clear glass bottle, Supelco) in chemically defined media under strict anaerobic conditions. The gas phase used for both strains was 80Vol.-% H_2_ in CO_2_ at 2.0 bar. *M. wolfeii* was grown at 60 °C and *M. smithii* at 37 °C, both shaking at 180 rpm.

### Methanogen media

M. wolfeii was grown in *Methanothermobacter marburgensis* medium as described in ^34^. *M. smithii* was grown in adapted DSMZ 141 Methanogen medium that contained 0.17 g KCl, 2 g MgCl_2_·6H_2_O, 0.125 g NH_4_Cl, 0.053 g CaCl_2_, 0.055 g KH_2_PO_4_, 0.84 g MgSO_4_, 3 g NaCl, 0.5 g Naacetate, 1 g yeast extract, 5 mL of trace element solution, 1 mL of FeII(NH_4_)_2_(SO_4_)_2_·6H_2_O solution, 50 µL of Na-resazurin solution (0.7 mg mL^−1^), and was filled up with 480 mL of ddH_2_O. The medium was brought to boil and was boiled for 5 min. After the temperature descended to 50 °C, 5 mL of vitamin solution, 2.5 g NaHCO_3_, 0.25g L-Cystein-HCl and 0.1 g Na_2_S·H_2_O were added. The pH was adjusted to 7 with HCl and the medium was filled up with ddH_2_O to a final volume of 500 mL. The medium was flushed with CO_2_ and transferred with a glass syringe into 100 mL serum bottles. 50 mL of the medium were aliquoted into serum bottles and sealed with blue rubber stoppers (pretreated by boiling ten times for 30 min in fresh ddH_2_O; 20 mm, Bellco) and open-top aluminum caps (9.5 mm opening, Merck Group). The sealed serum bottles containing the medium were autoclaved for 20 min at 120 °C. Composition of trace element was: 1.5 g Nitrilotriacetic acid, 3 g MgSO_4_·7H_2_O, 0.585 g MnCl_4_·4H_2_O, 1 g NaCl, 0.1 g FeSO_4_·7H_2_O, 0.18 g CoSO_4_·7H_2_O, 0.1 g CaCl_2_·2H_2_O, 0.18 g ZnSO_4_·7H_2_O, 0.006 g CuSO_4_, 0.02 g KAl(SO_4_)_2_·12H_2_O, 0.01 g H_3_BO_3_, 0.01 g Na_2_MoO_4_·2H_2_O, 0.03 g NiCl_2_·6H_2_O, 0.3 mg Na_2_SeO_3_·5H_2_O, 0.4 mg Na_2_WO_4_·2H_2_O. First nitrilotriacetic acid was dissolved and pH was adjusted to 6.5 with KOH. Then minerals were added, medium was filled up with ddH_2_O to a final volume of 1000 mL and pH was adjusted to 7.0 with KOH. Composition of Vitamin solution was: 2 mg Biotin, 2 mg Folic acid, 10 mg Pyridoxine-HCl, 5 mg Thiamine-HCl, 5 mg Riboflavin, 5 mg Nicotinic acid, 5 mg D-Ca-pantothenate, 0.1 mg Vitamin B12, 5 mg p-Aminobenzoic acid, 5 mg Lipoic acid and filled up with ddH_2_O to a final volume of 1000 mL.

### Cloning of *M. wolfeii* PeiW for recombinant protein production in textitE. coli

Genomic DNA from *M. wolfeii* was obtained by extracting with a phenol-chloroform extraction procedure. The gene encoding for PeiW (psiM100p36) was amplified by PCR ^35^ and the sequence of amplified peiW was verified by Sanger sequencing (Eurofins Genomics, Ebersberg, Germany). The insert and the Novagen vector pET-15b (Merck Group, Darmstadt, Germany) were digested with NdeI (NEB, Ipswich, MA, USA) and XhoI (NEB, Ipswich, MA, USA) and ligation of digested peiW and pET-15b with Quick ligase™ (NEB, Ipswich, MA, USA) generated the plasmid pET-15b_peiW, which was transformed into |textitE. coli Top10. The successful transformation was confirmed by a colony PCR and by Sanger sequencing (Eurofins Genomics, Ebersberg, Germany). Plasmid DNA was obtained by extracting with a Miniprep Kit (Pure Yield™, Promega, Madison, WI, USA), and pET-15b_peiW was transformed into *E. coli* DE3 BL21AI (Life technologies, Van Allen Way Carlsbad, CA, USA). All plasmids and primers used in this study are listed in Supplementary table 2 and Supplementary table 3 respectively.

### Protein expression and purification of PeiW

The expression of recombinant PeiW was induced at OD600 = 0.6 by the addition of 0.2 % (w/v) L-arabinose and 1 mmol L^−1^ IPTG to LB medium (100 µg mL^−1^ ampicillin). Cell cultures (3x 250 mL) were harvested after 3 h of incubation by a centrifugation of 15 min at 3170 g at 4 °C. To verify the expression of recombinant PeiW cell pellets were resuspended to a concentration of 0.01 optical density unit (ODU) µL^−1^ in 5x Laemmli buffer and 1 µL DTT (1 mol L^−1^) and lysed 5 min at 95 °C and 1050 rpm. Crude extracts were centrifuged 30 min at 16100 g at 4 °C and supernatant was loaded on two 12.5 % (w/v) polyacrylamide gels. The gel was stained with Commassie Brilliant Blue (50 % (v/v) ddH_2_O, 40 % (v/v) ethanol (ethanol absolute), 10 % (v/v) 100 Vol.-% acetic acid, 0.1 % (w/v) R-250 Brilliant Blue). For protein purification frozen cell pellets were resuspended in Ni-NTA lysis buffer (50 mmol L^−1^ NaH_2_PO_4_, 300 mmol L^−1^ NaCl, 10 mmol L^−1^ imidazole, at pH 8) at 4 °C and lysed by sonication. The lysate was centrifuged for 20 min at 14.000 rpm at 4 °C. The cleared lysate was incubated for 2 h at 4 °C with Ni-NTA agarose resin (Invitrogen) under slight agitation. The lysate with the beads was passed through a gravity flow chromatography column (Econo-Pac chromatography columns, Biorad). The column was washed with four column volumes of washing buffer (50 mmol L^−1^ NaH_2_PO_4_, 300 mmol L^−1^ NaCl, 20 mmol L^−1^ imidazole, at pH 8). Histagged proteins were eluted with 3x 1 mL elution buffer (50 mmol L^−1^ NaH_2_PO_4_, 300 mmol L^−1^ NaCl, 250 mmol L^−1^ imidazole, pH = 8) and fractions were collected separately. The fractions containing the protein of interest were loaded on a PD-midi Trap G25 column (GE Health Care) and eluted with 1.5 mL of 50 mmol L^−1^ Hepes buffer (at pH 7) for buffer exchange.

### Cloning of *M. smithii* SepF and FtsZ for recombinant protein production in *E. coli*

Genomic DNA from *M. smithii* was obtained by resuspending a pellet of densely grown culture in ddH_2_O and boiling it for 5 min at 99 °C. Subsequently, 1 µL of archaeal suspension was used as template in each 50 µL PCR reaction to amplify the genes encoding for *M. smithii ftsZ* (MSM_RS03130), *sepF* full length (SepF_FL_, MSM_RS02010) as well as only the core region of *sepF* (SepF_core_) comprising amino acids from 54 to 149. The products of the right size were cloned in to the pT7 vector containing an N-terminal 6xHis-SUMO tag by Gibson assembly. The constructs were transformed into chemically competent DH5α*E. coli* cells. The successful transformation was confirmed by a colony PCR and by Sanger sequencing (Eurofins Genomics, France).

### Protein expression and purification

Protein expression and purification was carried out as previously describes in ^19^. In brief, N-terminal 6xHis-SUMO-tagged SepF_FL_ and SepF_core_ from *M. smithii* were expressed in *E. coli* BL21 (DE3). After 4 h at 37 °C cells were grown for 24 h at 20 °C in 2YT complemented auto-induction medium ^36^ containing 50 µg mL^−1^ kanamycin. Cell pellets were resuspended in 50 mL lysis buffer (50 mmol L^−1^ Hepes pH8, 300 mmol L^−1^ NaCl, 5 % (v/v) glycerol, 1 mmol L^−1^ MgCl2, benzonase, lysozyme, 0.25 mmol L^−1^ TCEP, EDTA-free protease inhibitor cocktails (ROCHE)) at 4 °C and lysed by sonication. The lysate was centrifuged for 30 min at 30.000x g at 4 °C and loaded onto a Ni-NTA affinity chromatography column (HisTrap FF crude, GE Healthcare). His-tagged proteins were eluted with a linear gradient of buffer B (50 mmol L^−1^ Hepes at pH 8, 300 mmol L^−1^ NaCl, 5 % (v/v) glycerol, 1 mol L^−1^ imidazole). The eluted protein of interest was dialysed at 4 °C overnight in SEC buffer (50 mmol L^−1^ Hepes at pH 8, 150 mmol L^−1^ NaCl, 5 % (v/v) glycerol) in the presence of the SUMO protease (ratio used, 1:100). The cleaved protein was concentrated and loaded onto a Superdex 75 16/60 size exclusion (SEC) column (GE Healthcare) pre-equilibrated at 4 °C in 50 mmol L^−1^ Hepes at pH 8, 150 mmol L^−1^ NaCl, 5 % (v/v) glycerol. The purified protein was concentrated, aliquoted, flash frozen in liquid nitrogen and stored at −80 °C. N-terminal 6xHisSUMO-tagged *M. smithii* FtsZ was produced and purified as described above, except induction was performed at 37 °C, KCl was used instead of NaCl in all the buffers and a TALON FF crude column (GE Healthcare) was used for affinity chromatography.

### Thermal shift assay

Tests were performed using 96 well plate and 20 µL of reaction volume per well. A total of 3 µg of *Ms*SepF_core_ was used per assay in a buffer containing 150 mmol L^−1^ NaCl, 25 mmol L^−1^ Hepes at pH 8 and 5 % (v/v) glycerol. When used, FtsZ_CTD_ peptide (NEDQLDDFIDGIF purchased from Genosphere) was added to a final concentration of 1 mmol L^−1^. Next, 0.6 µL of 50X Sypro Orange solution (Invitrogen) was added to each well and samples were heated from 25 to 95 °C in 1 °C steps of 1 min each in a CFX96 Touch™ Real-Time PCR Detection System (BioRad). Excitation/emission filters of 492 and 516 nm were used to monitor the fluorescence increase resulting from binding of the Sypro Orange to exposed hydrophobic regions of the unfolding MsSepFcore. The midpoint of the protein unfolding transition was defined as the melting temperature (Tm).

### Crystallization

Crystallization screens were set up for *Ms*SepF_core_ apo form and in complex with tsZ_CTD_ using the sitting-drop vapour diffusion method at 18 °C in a Mosquito nanoliter-dispensing crystallization robot (TTP Labtech, Melbourn, UK) as detailed in ^37^. Optimal crystals of *Ms*SepF_core_ (14 mg mL^−1^) appeared after 5 days in a buffer containing 0.1 mol L^−1^ TRIS at pH 8.5 and 30 % (w/v) PEG 10K. The complex *Ms*SepF_core_-FtsZ_CTD_ crystallized in a buffer containing 0.2 mol L^−1^ (NH4)2SO4 and 30 % (w/v) PEG 8K after 1 week using *Ms*SepF_core_ at 14 mg mL^−1^ and FtsZ_CTD_ at 6 mg mL^−1^. Crystals were cryo-protected in the crystallization buffer containing 33 % (v/v) glycerol.

### Data collection, structure determination and refinement

X-ray diffraction data were collected at 100K using beamlines Proxima^−1^ and Proxima-2 at the Soleil synchrotron (GIF-sur-YVETTE, France). Both data sets were processed using XDS 38 and AIMLESS from the CCP4 suite ^39^. The crystal structures were solved by molecular replacement using Phaser ^40^ and a single monomer of the PDB 3ZIE as a search model. All structures were refined through several cycles of manual building with COOT ^41^ and reciprocal space refinement with BUSTER ^42^. The Supplementary table 1 shows the crystallographic statistics. Structural figures were generated using Chimera 43. Atomic coordinates and structure factors can be found in the protein data bank under the accession codes 7AL1 and 7AL2.

### Small unilamellar vesicles (SUVs) preparation

SUVs were prepared as previously describes in ^19^. In brief, reverse phase evaporation was used. A 10 mmol L^−1^ lipids chloroform solution was prepared and chloroform was removed by evaporation under vacuum conditions. The dried phospholipid film was resuspended in a mixture of diethyl ether and 25 mmol L^−1^ Hepes buffer at pH 7.4 and remaining diethyl ether was eliminated by reverse phase evaporation. Finally, SUVs were obtained by sonication during 30 min at 4 °C.

### Lipid peptide interaction (tryptophan fluorescence emission titration)

To estimate the partition coefficient (K_x_) between SUVs and SepF a similar protocol as described in ^19^ was used. In brief, we used the synthetic peptide WMGFTDALKRSLGF (purchased from Genosphere), which contains the SepF_M_ sequence of *M. smithii* with an extra W residue at the Nterminal. K_x_ is defined as the ratio of peptide concentration in the lipid and in the buffer phases. K_x_ can be expressed by the following equation:

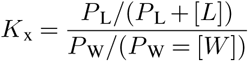

in which P_W_ represent the concentration of soluble peptide (in aqueous phase) and P_L_ the peptide concentration bound to lipid membranes (lipidic phase). [L] refers to the lipid concentration and [W] refers to the water concentration. K_x_ is directly related to the apparent dissociation constant as K_x_ * Kd = [W] with Kd * P_L_ = P_W_ * [L]. The KaleidaGraph software was used to fit the K_x_ to the experimental data. We used a FP8200 (Jasco, Tokyo, Japan) spectrophotometer equipped with a thermostatic Peltier ETC-272T at 25 °C. Experiments were performed in a high precision cell cuvette made of Quartz with a light Path of 10x 4 mm (Hellma, France). A bandwidth of 5 nm was used for emission and excitation. 1 µmol L^−1^ of peptide was used in Titration buffer (100 mmol L^−1^ KCl and 25 mmol L^−1^ Hepes at pH 7.4). We measured the florescence emission between 300 and 400 nm at a scan rate of 125 nm min^−1^ with an excitation wavelength of 280 nm. The obtained spectra were corrected by blank subtraction (SUV light scattering in titration buffer). Next, the maximum wavelength value (λ_max_) was calculated to measure the partition coefficient (K_x_).

### Electron microscopy

For negative stain sample preparations, incubations were performed at room temperature. SUVs (100 µmol L^−1^) and SepF_FL_ (50 µmol L^−1^) with or without FtsZ_CTD_ (100 µmol L^−1^) were incubated in a buffer containing 100 mmol L^−1^ KCl, 10 mmol L^−1^ MgCl_2_ and 25 mmol L^−1^ Pipes at pH 6.9 for 10 min. In order to visualize FtsZ filaments, FtsZ (30 µmol L^−1^) was incubated with or without SepF (20 µmol L^−1^) in polymerization buffer (2 mol L^−1^ KCl, 50 mmol L^−1^ Hepes at pH 7.4 and 10 mmol L^−1^ MgCl_2_) supplemented with freshly prepared 0.6 mol L^−1^ Trimethylamine N-oxide (TMAO) and 3 mmol L^−1^ GTP. Reactions were incubated for 10 min before applied onto the grid. For all samples, 400 mesh carbon coated grids (Electron Microscopy Sciences; CF 400-Cu) were glow-discharged on an ELMO system for 30 sec at 2 mA. 5 µL of sample was applied onto the grid and incubated for 30 s, the sample was blotted, washed in three drops of buffer (100 mmol L^−1^ KCl, 10 mmol L^−1^ MgCl_2_ and 25 mmol L^−1^ Pipes at pH 6.9) and then stained with 2 % (w/v) uranyl acetate. Images were recorded on a Gatan UltraScan4000 CCD camera (Gatan) on a Tecnai T12 BioTWINLaB6 electron microscope operating at a voltage of 120 kV.

### Statistical analysis of FtsZ filament diameter

Electron microscopy images of negatively stained FtsZ filaments (FtsZ+GTP) and FtsZ bundles (FtsZ+GTP+SepF) were analyzed using the public domain program Fiji. The diameter of each 130 FtsZ filaments and FtsZ bundles was measured in several images each and the data were extracted into Microsoft Excel 2016. Boxplots were created by using Microsoft Excel 2016 with the Real Statistics Resource Pack for Excel 2016 (http://www.real-statistics.com). Each data set for diameter of FtsZ filaments and FtsZ bundles was tested for normal distribution using the Kolmogornov-Smirnov test (KS-test). A 2-sample t-test was conducted to test whether the means of the diameter of FtsZ filaments and FtsZ bundles are significantly different. The statistical analysis was performed using DataLab version 4.0 (Epina GmbH, Pressbaum, Austria).

### Western blots and SepF and FtsZ quantification

Purified *Ms*SepF_core_ protein was used to raise antibodies in guinea pig (Covalab). Two peptides (CENAENGLEKLKSAADT and CGESDSGDRALESVHE) in equimolar amounts were used to raise antibodies in rabbit against *M. smithii* FtsZ (Genosphere Biotech). To prepare cell extracts, archaeal cell pellets were resuspended in PBS buffer (137 mmol L^−1^ NaCl, 10 mmol L^−1^ Na_2_HPO_4_, 1.8 mmol L^−1^ KH_2_PO_4_, 2.7 mmol L^−1^ KCl, at pH 7.4) together with LDS sample buffer (Invitrogen) and 100 mmol L^−1^ DTT, and disrupted at RT with 0.1 mm glass beads and using the MP FastPrep-24^TM^ 5G homogenizer. In order to test the expression of the proteins and quantify the amount of SepF and FtsZ in cells, crude extracts and purified recombinant SepF and FtsZ in serial dilution were loaded on a 4−12 % (w/v) BisTris PAGE gel (Invitrogen). The amounts of crude extract were 120 µg for FtsZ and 290 µg for SepF. Purified FtsZ protein was serially diluted from 100 ng to 6.26 ng, and purified SepF protein from 6.25 ng to 0.1 ng. After separation of proteins on SDS-PAGE gel, they were electro-transferred onto a 0.2 µm Nitrocellulose membrane. Membranes were blocked with 5 % (w/v) skimmed milk for 45 min at room temperature (RT). Membranes were incubated with either an anti-MsSepF antibody or an anti-MsFtsZ antibody, both at 1:500 dilution for 1 h at RT. After washing in TBS-Tween buffer (10 mmol L^−1^ Tris-HCl at pH 8; 150 mmol L^−1^ NaCl; Tween 20 0.05 % (v/v), the membranes were incubated with an anti-rabbit horseradish peroxidase-linked antibody for FtsZ and an anti-guinea pig horseradish peroxidase-linked antibody for SepF (Invitrogen), both at 1:5000 dilution for 45 min. The membranes were washed and revealed with HRP substrate (Immobilon Forte, Millipore) and imaged using the ChemiDoc MP Imaging System (BIORAD). Quantification of the bands was performed using the software Image Lab (Biorad) and the band volume was plotted against the ng loaded in order to obtain a strand curve. All uncropped blots are shown in Supplementary figure 2.

### Immunostaining

Cells of *M. smithii* grown to early exponential phase were harvested by 5 min of centrifugation at 3.5x g (all centrifugation steps were performed at 3.5x g). Pellets were washed in PBS buffer, pelleted, fix with 100 % ice cold methanol and stored at −20 °C. Cells were rehydrated and washed in 50 mmol L^−1^ HEPES buffer at pH 7 for 10 min at RT, followed by permeabilization of the archaeal pPG by incubation for 10 min with 3.5 µg of purified 6xHis-PeiW in 50 mmol L^−1^ HEPES buffer containing 1 mmol L^−1^ DTT at 71 °C. After the incubation, cells were allowed to cool on ice for 2 min. Cells were pelleted and washed one time in 50 mmol L^−1^ HEPES buffer and two times in PBS-T (PBS buffer with Tween 20 at 0.1 % (v/v)). Blocking was carried out for 1 h in PBS-T containing 2 % (w/v) bovine serum albumin (blocking solution) at RT. Cells were incubated with a 1:200 dilution of rabbit polyclonal anti-*Ms*FtsZ peptide antibody and 1:200 dilution of guinea pig polyclonal anti-*Ms*SepF antibody overnight at 4 °C in blocking solution. Upon incubation with primary antibody samples were pelleted and washed three times in PBS-T, and incubated with a 1:500 dilution of secondary Alexa555-conjugated anti-rabbit for FtsZ and Alexa488-conjugated anti-guinea pig for SepF (Invitrogen) in blocking solution for 1 h at RT. Unbound secondary antibody was removed by three washing steps in PBS-T. Finally, cells were resuspended in few µL of PBS and slides were prepared either for super resolution microscopy or epifluorescence microscopy (see below).

### Three-Dimensional Structured Illumination Microscopy (3D SIM) imaging and analysis

Archaeal cell suspensions were applied on high precision coverslips (No. 1.5H, Sigma-Aldrich) coated with 0.01 % (w/v) of Poly-L-Lysin. After letting the cells attach onto the surface of the coverslip for 10 min, residual liquid was removed, 8 µL of antifade mounting medium (Vectashield) were applied and the coverslip was sealed to a slide. SIM was performed on a Zeiss LSM 780 Elyra PS1 microscope (Carl Zeiss, Germany) using C Plan-Apochromat 63× / 1.4 oil objective with a 1.518 refractive index oil (Carl Zeiss, Germany). The samples were excited with laser at 488 nm and 561 nm and the emission was detected through emission filter BP 495-575 + LP 750 and BP 570-650 + LP 750, respectively. The fluorescence signal is detected on an EMCCD Andor Ixon 887 1K camera. Raw images are composed of fifteen images per plane per channel (five phases, three angles), and acquired with a Z-distance of 0.10 µm. Acquisition parameters were adapted from one image to one other to optimize the signal to noise ratio. SIM images were processed with ZEN software (Carl Zeiss, Germany) and then corrected for chromatic aberration using 100 nm TetraSpeck microspheres (ThermoFisher Scientific) embedded in the same mounting media as the sample. For further image analysis of SIM image z stacks we used Fiji (ImageJ) Version 2.0.0-rc-68/1.52i. Namely, we assigned a color to the fluorescent channel, stacks were fused to a single image (z projection, maximum intensity), stacks were rotated 90° (resliced) prior z projection for the side view, and videos were created via 3D projection. Regions of interest were cut out and, for uniformity, placed on a black squared background. Figures were compiled using Adobe Photoshop 2020 and Illustrator 2020 (Adobe Systems Inc. USA).

### Morphometric and fluorescence measurements

1 µL of immunolabelled *M. smithii* cells solution together with 1 µL of Vectashield (Vector Labs) were applied to an 1 % (w/v) agarose covered microscopy slide and imaged using a Zeiss Axioplan 2 microscope equipped with an Axiocam 503 mono camera (Carl Zeiss, Germany). Epifluorescence images were acquired using the ZEN lite software (Carl Zeiss, Germany) and processed using the public domain program Fiji (ImageJ) Version 2.0.0-rc-68/1.52i in combination with plugin MicrobeJ Version 5.13l. The cell outlines were traced and cell length, width fluorescence intensity along the cell length and presence of fluorescent maxima were measured automatically. Automatic cell recognition was manually double-checked. For the mean fluorescence intensity plots cells were automatically grouped into four classes according to the detected FtsZ (0-3 fluorescent maxima detected) and the corresponding FtsZ and SepF mean fluorescence intensity of each group was plotted against the normalized cell length. For the plots showing the relative position of detected maxima, cells were automatically grouped into four classes for SepF (1-4 fluorescent maxima detected) and three for FtsZ (1-3 fluorescent maxima detected). Both, fluorescence intensity and relative position of detected maxima plots were created in MicrobeJ, representative cells were chosen from bigger fields of few and their brightness and contrast were adapted in Fiji. Figures were compiled using and Illustrator 2020 (Adobe Systems Inc. USA).

### Sequence analysis

FtsZ sequences from either Archaea or Bacteria were aligned with MAFFT using the L-INS-i algorithm ^44^. Only the Cterminal domain of FtsZ was kept and after several cycles of trimming and realigning, the aligned FtsZ_CTD_ was uploaded to https://weblogo.berkeley.edu/logo.cgi to create the WebLogo ^45^. For structure-based SepF alignment, representative sequences from Bacteria and Archaea were manually chosen. Sequences where aligned using T-Coffee ^46^. Graphical representation was made using ENDscript server ^47^. For secondary structure prediction, sequences were aligned using MAFFT with the L-INS-i algorithm ^44^ and secondary structures were detected using Ali2D ^48^. Figures were compiled using Illustrator 2020 (Adobe Systems Inc. USA).

### Distribution, phylogeny, and synteny analysis

A customized local genome database was compiled for 190 Bacteria and 150 Archaea representative of all major phyla in NCBI (as of January 2020). The presence of FtsZ, FtsA and SepF homologues was assessed by performing HMM-based homology searches using custom made HMM models and the HMMER package ^49, 50^ (default parameters). Separate searches were carried out with archaeal and bacterial queries and absences were checked with TBLASTN ^51^ or individual HMMER searches against all available taxa in NCBI corresponding to those phyla. Results were mapped onto the bacterial and archaeal reference phylogenies using iTOL ^52^. For phylogenetic analysis, FtsZ, FtsA and SepF homologues were aligned using MAFFT (L-INS-i algorithm) ^53^. Columns with excess gaps (>80 % for FtsA/SepF, >70 % for FtsZ) were removed with trimAl ^54^. Maximum likelihood trees were generated using IQ-TREE v1.6.7.2 ^55^, using the SH-aLRT branch test (‘-alrt 1000’ option) and the ultrafast bootstrap approximation ^56^ (option ‘–bb 1000’) for branch supports. Exchangeability matrices and overall evolutionary models for each alignment were determined using ModelFinder ^57^, with heterogeneity rates modeled with the gamma distribution (four rate categories, G4), or free-rate models (+R5 to +R9, depending on the protein). For each archaeal genome, ten genes upstream and downstream ftsZ1 and ftsZ2 and sepF were extracted and the corresponding proteins were subjected to all-vs-all pairwise comparisons using BLASTP v2.6.0 ^51^ with default parameters. From the output of the BLASTP search, protein families were assembled with SILIX v1.2.951 ^58^ with default parameters. HMM profiles were created for the families containing members of three or more archaeal lineages using the HMMER package ^50^. The families were annotated manually by using BLASTP v2.6.0 ^51^ and the Conserved Domain Database from NCBI ^59^. MacSyFinder ^60^ was then used to identify, in each archaeal taxon, clusters containing at least three genes with a separation no greater than five other genes.

## Acknowledgments

This work was partially supported by grants from the Institut Pasteur (Paris), the CNRS (France) and the Agence Nationale de la Recherche (PhoCellDiv, ANR-18-CE11-0017-01 and ArchEvol, ANR-16-CE02-0005-01). N.P. is supported by a Pasteur-Roux Postdoctoral Fellowship from the Institut Pasteur (Paris). A.S. and D.M. are part of the Pasteur Paris University (PPU) International PhD Program, funded by the European Union’s Horizon 2020 research and innovation program under the Marie Sklodowska-Curie grant agreement No 665807. M.G. acknowledges support from Programa de Desarrollo de las Ciencias Básicas and Agencia Nacional de Investigación e Innovación, Uruguay. We thank A. Chenal for help with the lipid interaction studies. We gratefully acknowledge the core facilities at the Institut Pasteur C2RT, A. Haouz, P. Weber, C. Pissis (PFC). We thank the staff of the synchrotron SOLEIL for assistance and support in using beamlines PX1 and PX2. We thank A. Salle for help with the 3D SIM. We gratefully acknowledge the UTechS Photonic BioImaging (Imagopole), C2RT, Institut Pasteur (Paris, France) as well as the France–BioImaging infrastructure network supported by the French National Research Agency (ANR-10–INSB–04; Investments for the Future), and the Région Ile-de-France (program Domaine d’Intérêt Majeur-Malinf) for the use of the Zeiss LSM 780 Elyra PS1 microscope. We acknowledge technical assistance by Barbara Reischl.

## Author contributions

N.P performed immunolabelling, epifluorescence microscope and 3D SIM imaging and image analysis. N.P. and A.S. conducted the protein biochemistry and purification for structural and biophysical studies. N.P. and A.S. carried out the biochemical and biophysical studies of protein-protein interactions. A.S. carried out binding studies of lipid membrane-protein interactions. D.M. and M.G. performed sequence and phylogenetic analyses. H.P. and S.K.-M.R.R. carried out PeiW purification and provided material. A.S. and P.M.A. carried out the crystallogenesis and crystallographic studies.

A.M.W. and A.S.R. performed the negative stain EM studies. N.P. and S.K.-M.R.R. performed statistical analysis. N.P., A.S. and D.M. made the figures. A.S., N.P., A.M.W., P.M.A and S.G. wrote the manuscript. All authors edited the final version of the manuscript.

