## Supplementary Information for "SepF is the FtsZ-anchor in Archaea: implications for cell division in the Last Universal Common Ancestor"

**Pende N., Sogues A., *et al*. 2020**

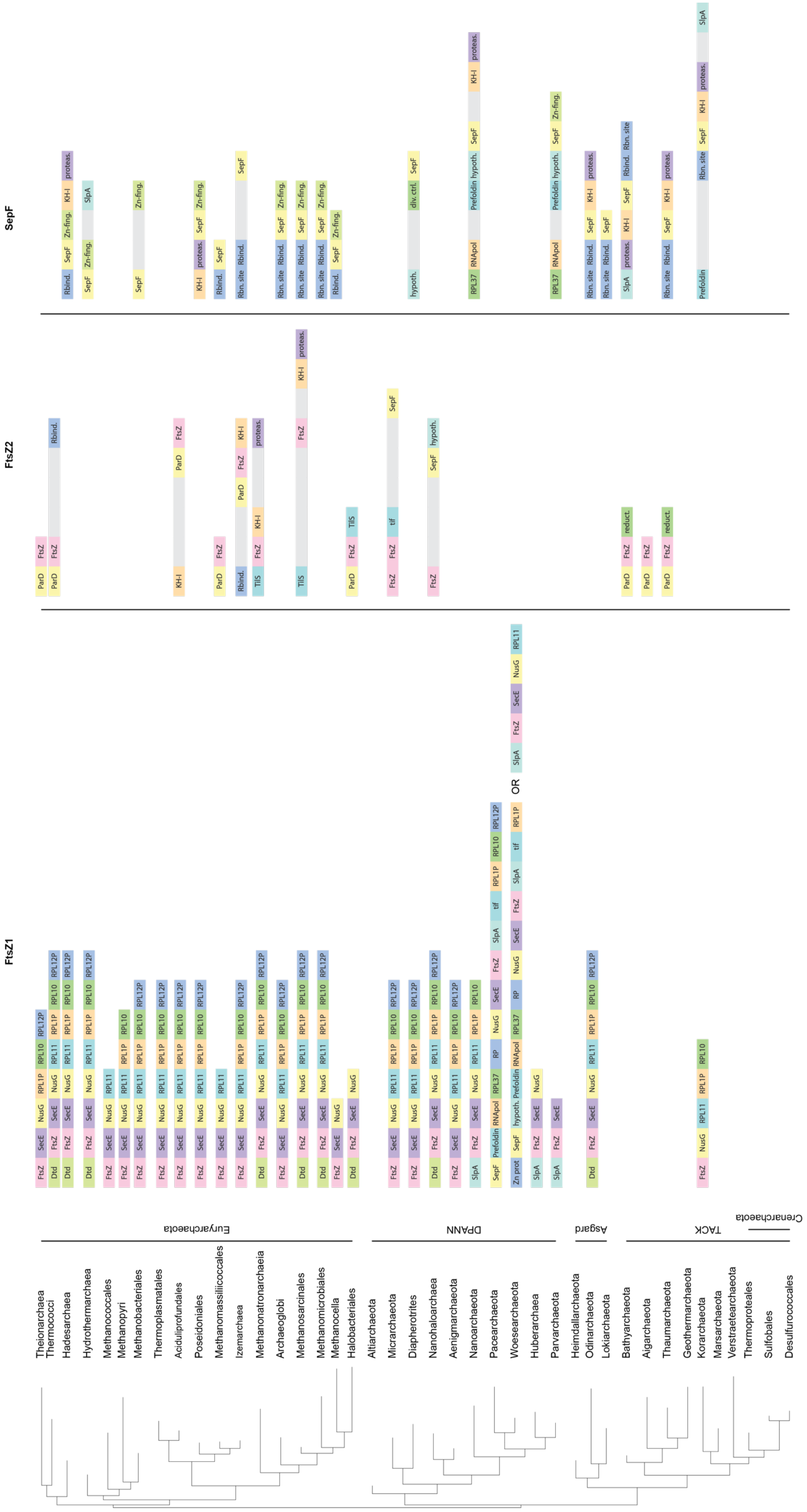


**Supplementary Figure 1. Genomic context of archaeal FtsZ1, FtsZ2 and SepF homologues mapped on a schematic reference phylogeny of the Archaea.** The genomic context of the gene coding for FtsZ1 is well conserved in most archaeal lineages. Most of the genes around *ftsZ1* are involved in transcription, translation and regulation. In contrast, the genomic contexts of the genes coding for SepF and FtsZ2 are less conserved. In some members of the DPANN superphylum *sepF* can be found in the same conserved cluster as *ftsZ1* of *ftsZ2*, supporting the functional link between the two proteins.

div. ctrl.: ORC1-type DNA replication protein; dtd: D-aminoacyl-tRNA deacylase; FtsZ: FtsZ, hypoth.: Uncharacterized protein family (UPF0147); KH-I: K homology RNA-binding domain type I; proteas.: Proteasome subunit; Rbind.: RNA-binding protein; Rbn.: site RNA binding site; reduct.: Nitro FMN reductase; RNApol: DNA-directed RNA polymerase; RP: 50S ribosomal protein; RPL10: Ribosomal protein L10 family; RPL11: 50S ribosomal protein L11; RPL12P: 50S ribosomal protein L12P; RPL1P: 50S ribosomal protein L1P; RPL37: 50S ribosomal protein L37; tif: translation initiation factor IF-5A; Zn-fing: ZPR1 zinc-finger domain protein. Not conserved genes are represented in grey.

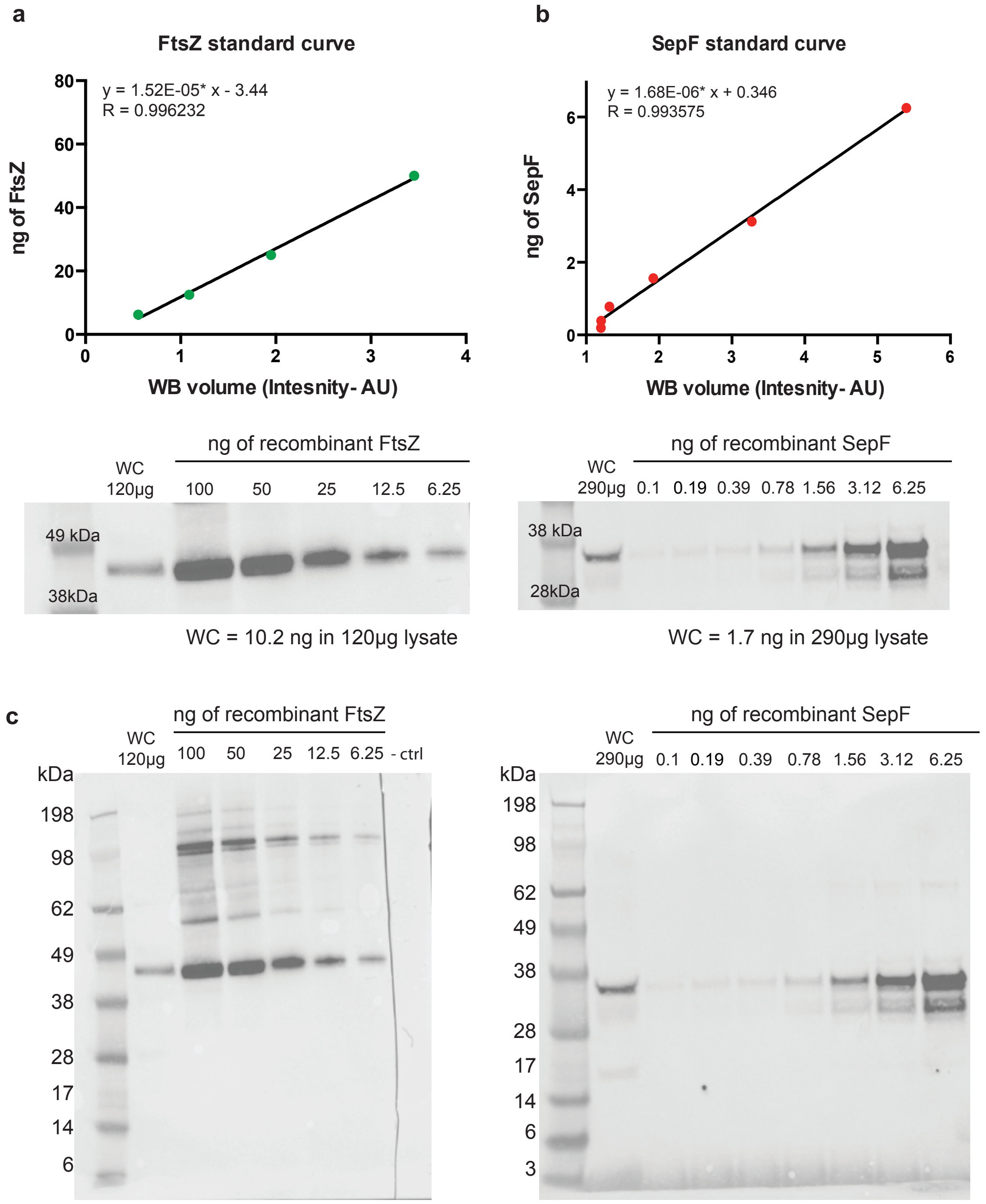


**Supplementary Figure 2. Ratio of FtsZ/SepF in *M. smithii* and characterization of the specific anti-*Ms*FtsZ and anti-*Ms*SepF antibodies. a.** Serial dilutions of recombinant FtsZ (100 ng to 6.25 ng). Band volumes were plotted against amount of FtsZ in order to calculate the linear regression. 120 µg of the whole cell extract in exponential phase were loaded and the total amount of FtsZ was calculated. **b.** Serial dilutions of recombinant SepF (6.25 ng to 0.1 ng). Band volumes were plotted against amount of SepF in order to calculate the linear regression. 290 µg of the whole cell extract in exponential phase were loaded and the total amount of SepF was calculated. **c.** Full uncropped Western Blots. Molecular weight markers (MW, in kDa) are shown on the side of the blot. The data shown here are representative for experiments performed at least twice.

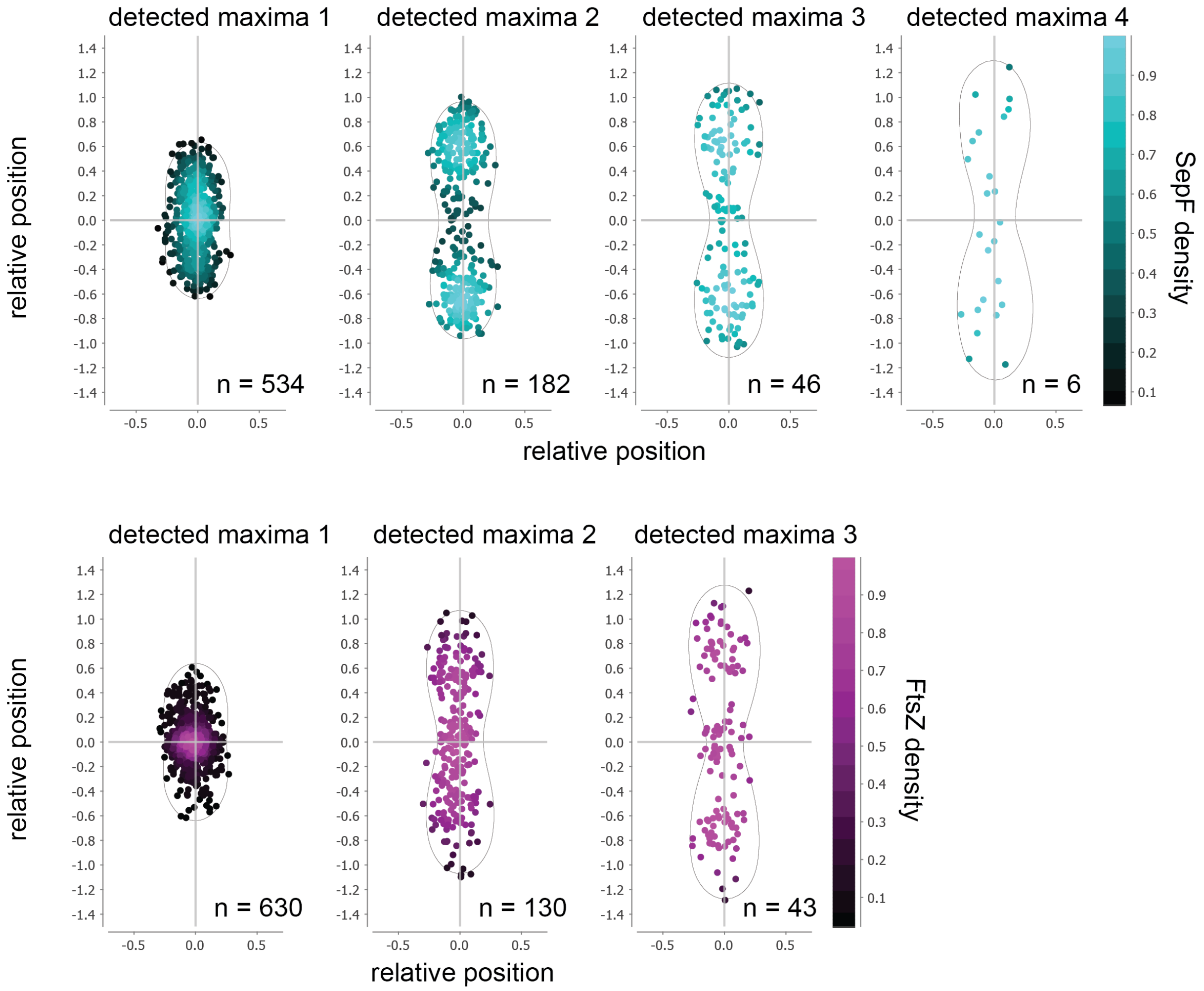


**Supplementary Figure 3. Localization of SepF and FtsZ maxima within *M. smithii* cells.** Relative position of detected fluorescent maxima within the cell grouped into four classes for SepF (maxima detected 1-4, upper panel) and three for FtsZ (maxima detected 1-3, lower panel). n, number of cells for each group. The data shown here are representative for experiments performed three times.

a

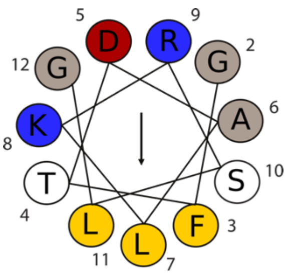


b

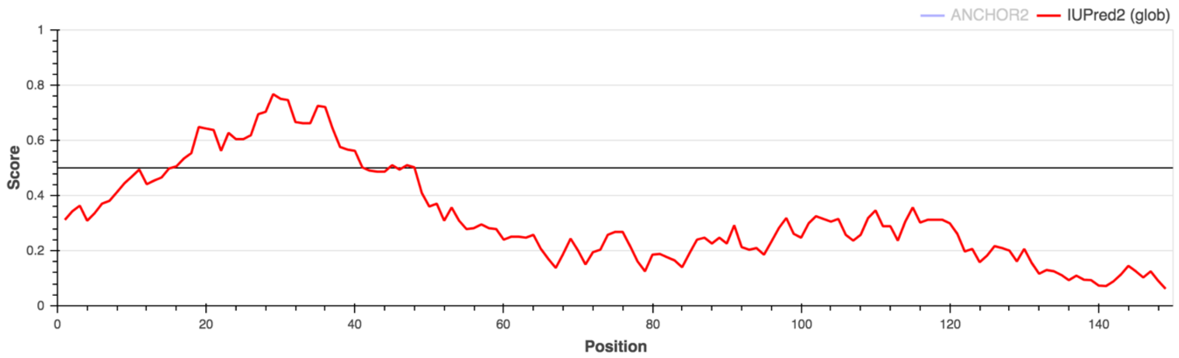


**Supplementary Figure 2**. **Functional and structural domains of SepF from *M. smithii.* a**. Predicted amphipathic helix for the M domain comprising amino acids from 2 to 12. Amino acids proprieties color coded: gray (non-charged) yellow (apolar), red (negatively charged) and blue (positively charged). Prediction was made using HELIQUEST [1]. **b**. Intrinsically disordered region prediction of SepF by IUPred2A software [2].

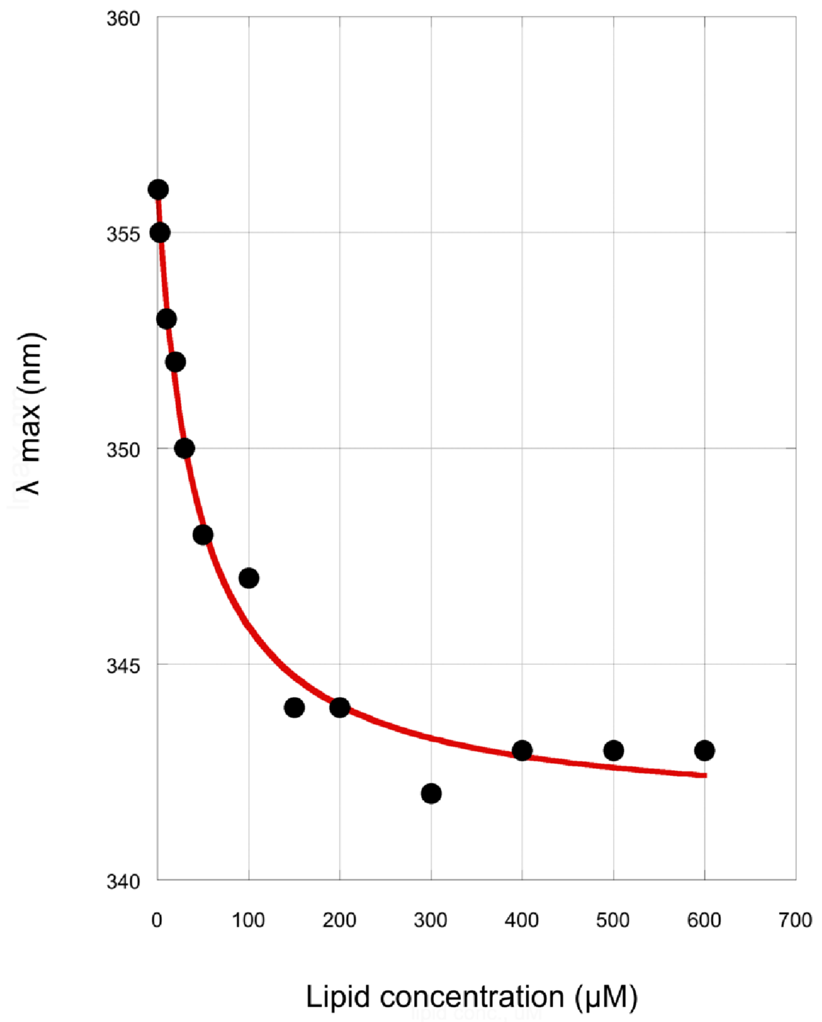


**Supplementary Figure 5. SepF$_M$ peptide – Membrane interaction.** Tryptophan fluorescence titration assay using a modified SepF$_M$ peptide (including a N-terminal tryptophan residue) as a function of lipid concentration (see Material and Methods for details).

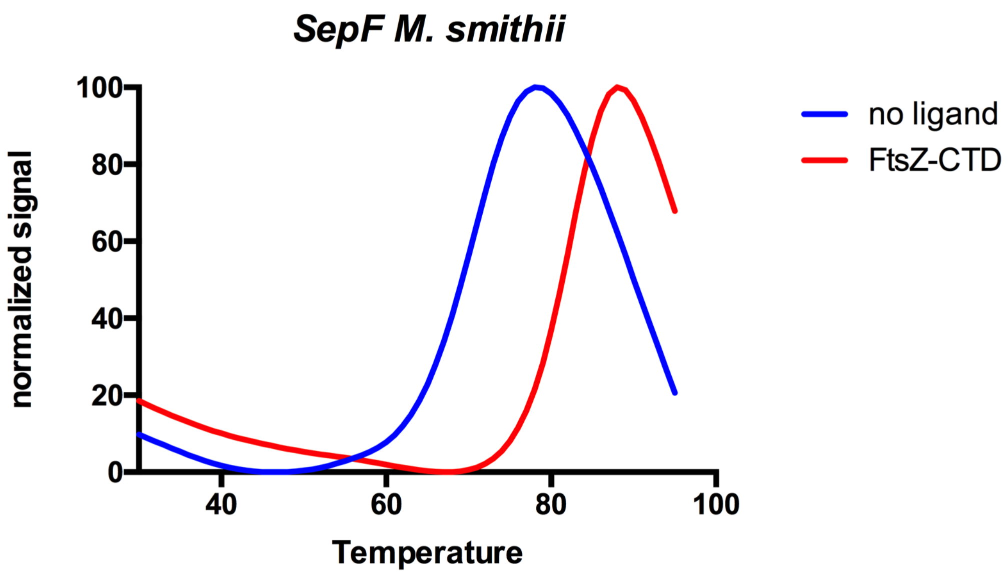


**Supplementary Figure 6. Thermal shift assays on *Ms*SepF.** Thermal denaturation curves of MsSepF alone (blue) and upon addition of the $FtsZ_{CTD}$ peptide (red). The significant increase of the protein thermostability (Tm values of 70.6 °C and 82.3 °C, respectively) indicates a strong interaction *in vitro*. Experiments were performed in triplicates.

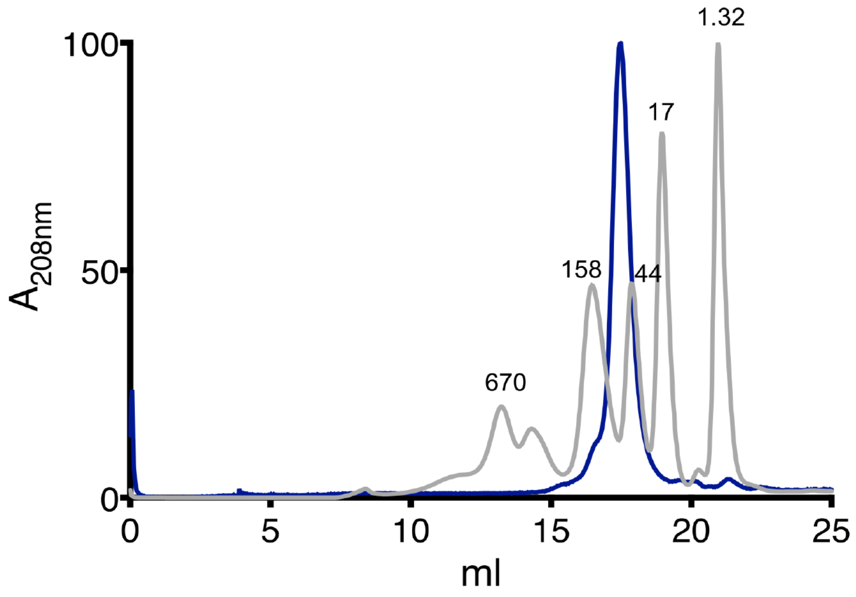


**Supplementary Figure 7. Size exclusion chromatography of the full-length *Ms*SepF shows the dimeric state of the protein in solution**. *Ms*SepF full-length (blue) on a Superdex S200 10/300 column. The molecular weight (MW) markers are shown in gray with corresponding MW indicated above. The data shown here are representative for experiments performed at least twice.

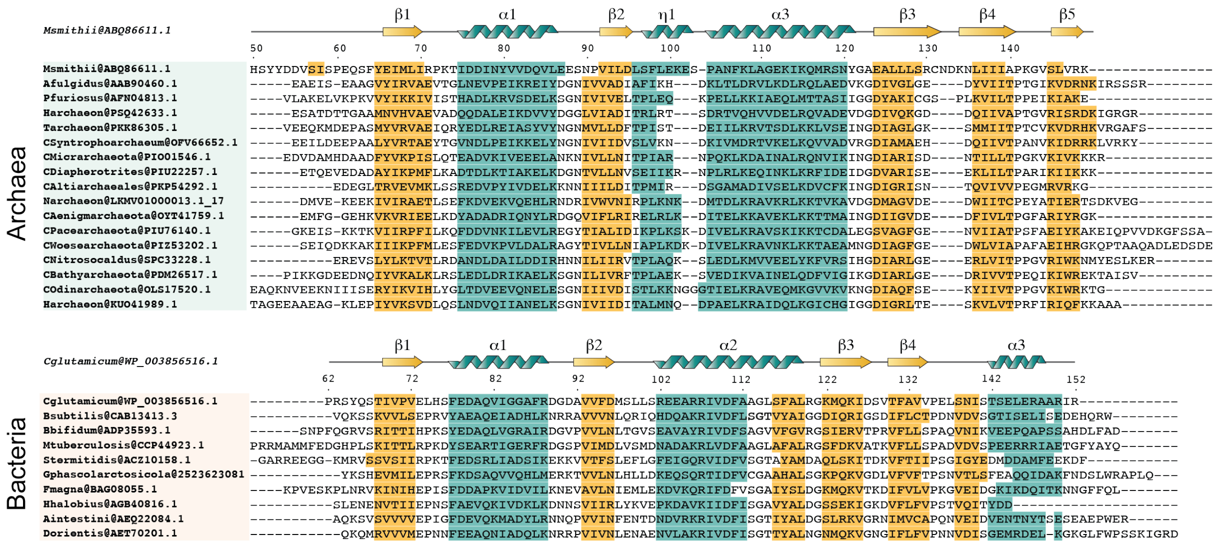


**Supplementary Figure 8**. **Prediction of the secondary structure of archaeal and bacterial SepF**. Sequences from diverse and representative linages in Archaea and Bacteria were chosen and aligned based on structural features. Secondary structural elements depicted above the sequence correspond to the structure of *M.smithii* and *C. glutamicum* (PDB: 6SCP). Turquoise indicates predicted α-helical structures whereas yellow indicates predicted β-strands. Prediction was performed using PSIPRED [3] and Ali2D [4].

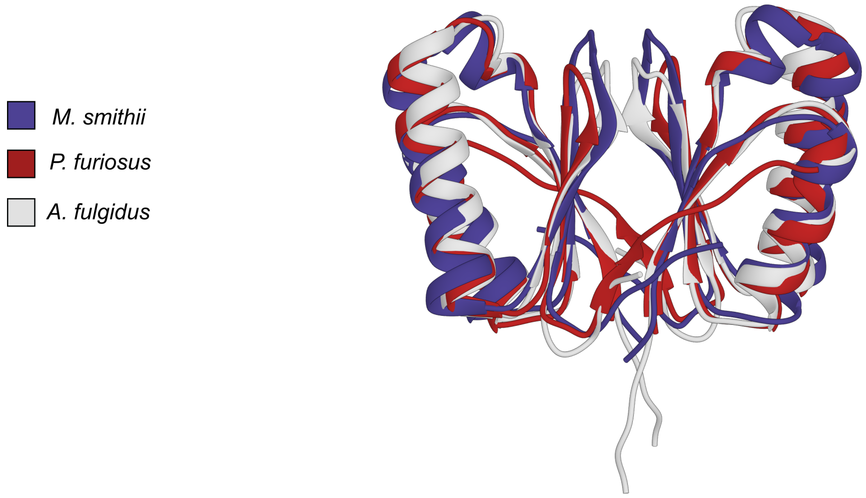


**Supplementary Figure 9**. **Comparison of available archaeal SepF structures**. Superposition of all the available archaeal SepF crystal structures (RMSDs of 1.5 - 1.9 Å for 75 – 77 aligned residues) show the same folding and dimer interface. The three structures contain the same secondary structural elements (α1 to α2, η1 and β1 to β5). *Archaeoglobus fulgidus* (PDB: 3ZIE), *Pyrococcus furiosus* (PDB: 3ZIG) [5].

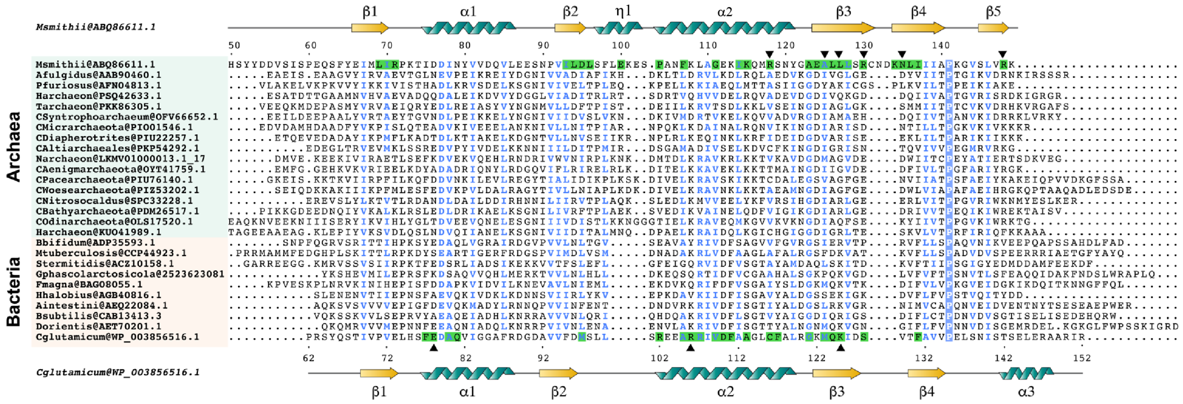

**Supplementary Figure 10. Archaeal and bacterial SepF structure-based alignment.** Sequences from diverse and representative linages in Archaea and Bacteria were chosen and aligned based on structural features. SepF from M.smithii and from *C. glutamicum* (PDB: 6SCP) were chosen as a structural models and their secondary structural elements are shown above. Conserved positions are indicated in blue; residues that participate in the $FtsZ_{CTD}$ interaction are highlighted in green and those that establish hydrogen bonds are marked with a black triangle above. Graphical representation was made using ENDscript server [6] and improved manually.

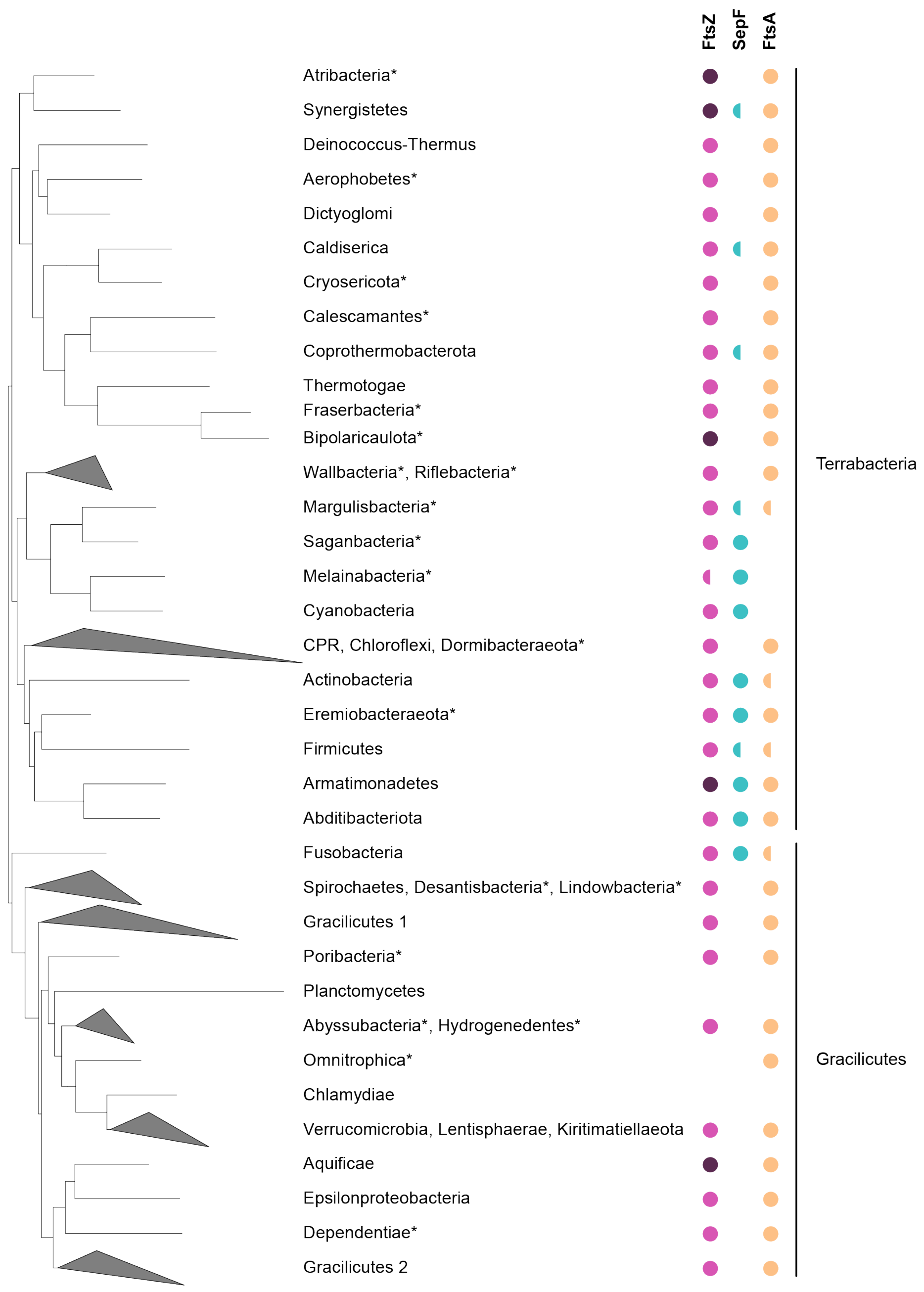


**Supplementary Figure 11. Distribution of FtsZ, SepF and FtsA on a schematic representation of the Bacteria phylum-level phylogeny.** FtsZ (magenta) is present in most bacterial phyla, with the exception of some phyla within the PVC superphylum (Planctomycetes, Omnitrophica and Chlamydiae), which are known to have specific FtsZ-less cytokinesis. Both SepF and FtsA are also absent in Planctomycetes and Chlamydiae. Two copies of FtsZ can be identified in Atribacteria, Synergistetes, Biopolaricaulota, Armatimonadetes and Aquificae (dark magenta). Only SepF is present in some phyla (Synergistetes, Caldiserica, Coprothermobacterota, Margulisbacteria, Melainabacteria, Cyanobacteria, Actinobacteria, Eremiobacteraeota, Firmicutes, Armatimonadetes, Abditibacteriota and Fusobacteria), most belonging to the Terrabacteria. In contrast, all

Gracilicutes except Fusobacteria, and the remaining Terrabacteria phyla have only FtsA. Finally, some members of Synergistetes, Caldiserica, Coprothermobacterota, Margulisbacteria, Actinobacteria, Eremiobacteraeota, Firmicutes, Armatimonadetes, Abditibacteriota and Fusobacteria have both FtsA and SepF. Semicircles indicate that the corresponding protein could not be identified in all the analyzed taxa in the linage displayed. * indicates uncultured Candidate phyla for which many genomes are incomplete. These phyla were not included in phylogenetic reconstructions. For full data see Supplementary data 2.

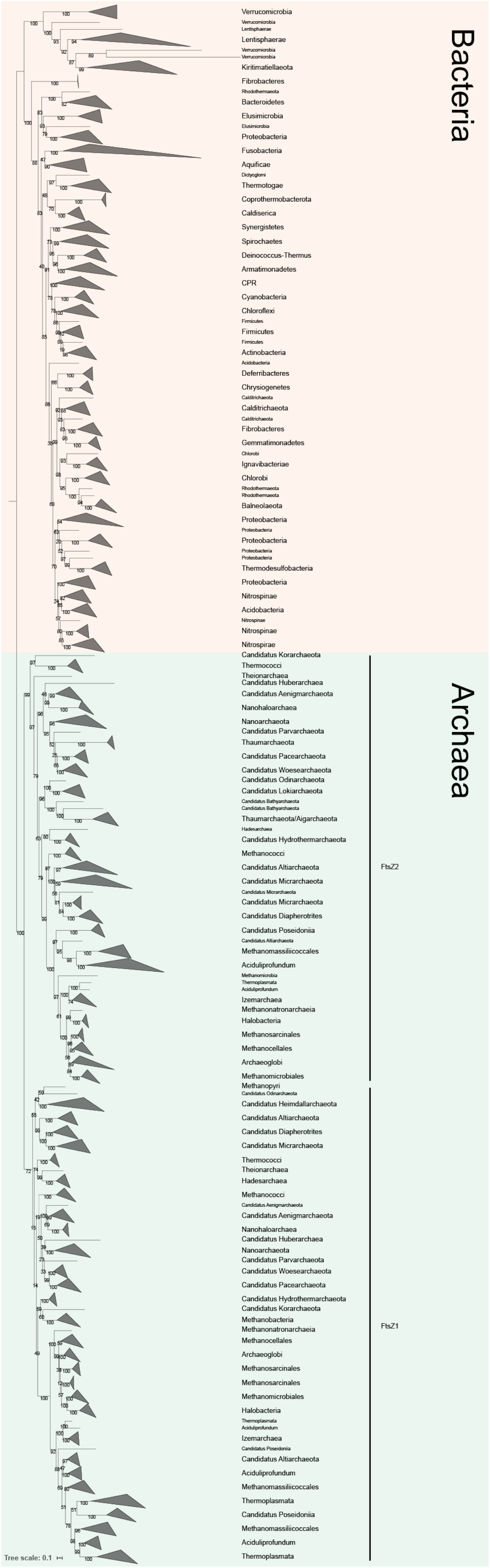
Bacteria
Archaea
FtsZ2
FtsZ1
Tree scale: 0.1

**Supplementary Figure 12. Phylogeny of FtsZ homologues in Bacteria and Archaea.** Maximum likelihood tree of FtsZ inferred with IQ-TREE v1.6.7.2 (ModelFinder best-fit model LG+R10) [7, 8] from an alignment of 429 sequences and 422 amino acid positions. Numbers at nodes represent ultrafast bootstrap supports [9]. The scale bar represents the average number of substitutions per site. The internal topologies roughly recapitulate known phylogenetic relationships, suggesting that FtsZ was already present in the LUCA. The separation of the two archaeal FtsZ1 and FtsZ2 copies suggests that they arose from an early gene duplication.

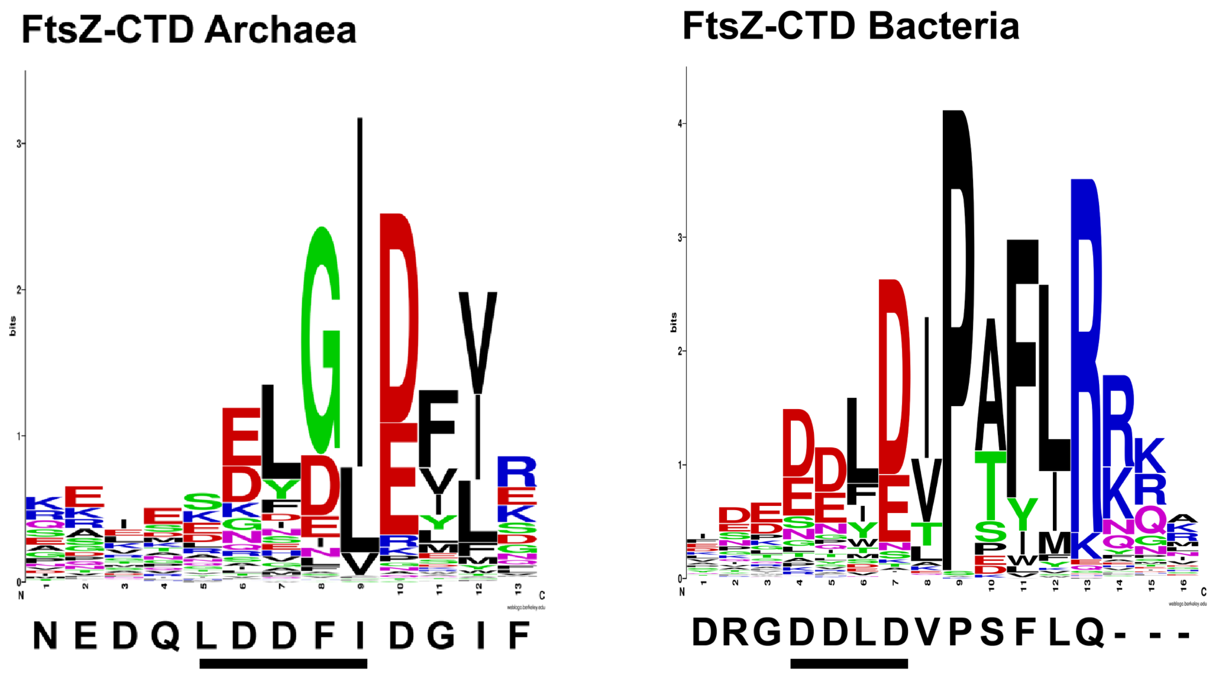


**Supplementary Figure 13. WebLogo plot of the FtsZ-CTD in archaea and bacteria.** For Archaea 330 sequences and 23 positions were taken whereas 266 and 29 positions were used for Bacteria. *M.smithii* (in Archaea) and *C. glutamicum* (in Bacteria) sequence are shown aligned below. Underlined amino acids represent the conserved positions that bind to a similar groove formed between $\alpha 2$ and $\beta 3$ in the SepF monomer. The conserved consensus motif of the archaeal $FtsZ_{CTD}$ is shorter than the bacterial (6 vs 10), the ordered $FtsZ_{CTD}$ peptide bound to *Ms*SepF has the same length as that seen in the bacterial complex, suggesting that a consensus length of about 10 amino acids is required for the SepF-FtsZ interaction.

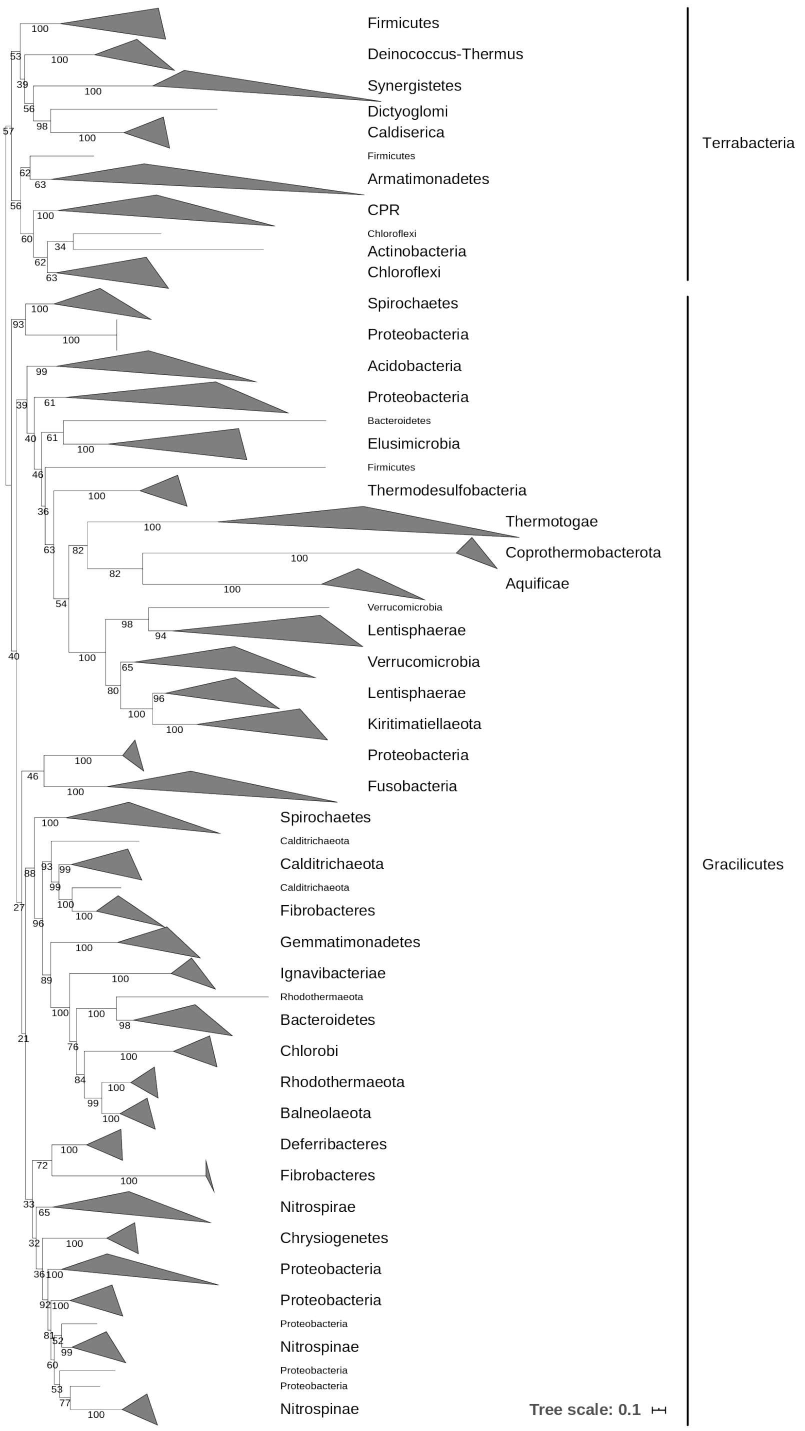


**Supplementary Figure 14. Phylogeny of FtsA homologues in Bacteria.**

Maximum likelihood tree of FtsA inferred with IQ-TREE v1.6.7.2 (ModelFinder best-fit model LG+R6) [7, 8] from an alignment of 163 sequences and 426 amino acid positions. Numbers at nodes represent ultrafast bootstrap supports [9]. The scale bar represents the average number of substitutions per site. The tree roughly recapitulates known phylogenetic relationships, suggesting that FtsA was already present in the LBCA (Last Bacterial Common Ancestor).

**Supplementary Table 1.** Crystallographic data.

| | ***Ms*SepF$_c$** | ***Ms*SepF$_c$ + FtsZ$_{CTD}$** |
|---|---|---|
| ***Data collection*** | | |
| Space group | P $3_1$ 2 1 | P $6_1$ 2 2 |
| Cell dimensions | | |
| a, b, c (Å) | 53.05, 53.05, 53.91 | 64.85, 64.85, 107.92 |
| α, β, γ (°) | 90, 90, 120 | 90, 90, 120 |
| Resolution (Å)* | 45.95–1.4 (1.42–1.4) | 38.91 - 2.7 (2.83 - 2.7) |
| $R_{sym}$ | 0.056 (1.029) | 0.076 (0.702) |
| I/σ(I) | 27.1 (3.4) | 15.4 (2.7) |
| Completeness (%) | 99.9 (100) | 95.7 (96.3) |
| Redundancy | 19.6 (19.9) | 5.5 (5.7) |
| ***Refinement*** | | |
| Resolution (Å) | 1.4 | 2.7 |
| Number of reflections | 17673 | 3806 |
| R-work/R-free | 0.182 / 0.216 | 0.229 / 0.255 |
| Number of atoms | | |
| protein | 684 | 746 |
| ligands/ions | 8 | - |
| water | 57 | - |
| B-factors (Å$^2$) | | |
| protein | 25.47 | 61.5 |
| ligands/ions | 25.49 | - |
| water | 37.47 | - |
| RMS deviations | | |
| Bond length (Å) | 0.009 | 0.008 |
| Bond angles (°) | 1.057 | 1.02 |
| ***PDB code*** | **7AL1** | **7AL2** |

* Values in parenthesis refer to the highest recorded resolution shell.

**Supplementary Table 2.** Bacterial strains and plasmids used in this study.

| Strain or plasmid | Characteristics | References |
|---|---|---|
| ***E. coli*** | | |
| **DH5*α*** | F- endA1 Φ80dlacZΔM15 Δ(lacZYA-argF)U169 recA1 relA1 hsdR17(rK–mK+) deoR supE44 thi-1 gyrA96 phoA λ–; strain used for general cloning procedures | [10] |
| Top10 | F- *mcrA* Δ( *mrr-hsd*RMS-*mcr*BC) Φ80*lac*ZΔM15 Δ *lac*X74 *rec*A1 *ara*D139 Δ( *araleu*)7697 *gal*U *gal*K *rps*L (StrR) *end*A1 *nup*G; strain used for general cloning procedures | |
| BL21(DE3) | F- ompT hsdSB(rB–mB–) gal dcm (DE3); host for protein production | [11] |
| ***M. wolfeii*** | | |
| DSM 2970 | Methanogenic, anaerobic, 60°C, type strain, | [12] [13] |
| ***M. smithii*** | | |
| DSM 861 | Methanogenic, anaerobic, 37°C, type strain | [14] |
| **Plasmids** | | |
| **pET-15b** | T7 promotor, His-tag, multiple cloning sites (*Nde*I - *Bam*H I), lacI, AmpR | |
| **pET-15b_*peiW*** | AmpR, pET derived for *M. wolfeii* peiW recombinant expression containing a N-terminal His-tag followed by a thrombin cleavage site | [15] |
| **pET-SUMO-*sepF*_full** | KanaR; pET derivate for *M.smithii* SepF recombinant expression containing a N-terminal His-tag followed by a SUMO protease cleavage site | This work |
| **pET-SUMO-*sepF*_core** | KanaR; pET derivate for *M.smithii* SepF (53-149) recombinant expression containing a N-terminal His-tag followed by a SUMO protease cleavage site | This work |

**Supplementary Table 3**. Oligonucleotides used in this study

| Oligonucleotide | Sequence (5'→ 3') and properties * |
|---|---|
| **Construction of pET-15b_*peiW*** | |
| PeiWF2 | AGGTGATCATATGGAAGTGGGGCTAAATG |
| PeiWR2 | AACAACTCGAGCATGTCTCTGCCACAAAC |
| **Construction of pET-SUMO-*sepF*_full** | |
| Ms_SepF_full_f | **CGAACAGATTGGTGGC**ATGGGTTTCACTGATG |
| Ms_SepF_full_r | **GTTAGCAGCCGGATCT**CTACTTTCTAACTAAACTGACTCC |
| **Construction of pET-SUMO-*sepF*_core** | |
| Ms_SepF_core_f | **CGAACAGATTGGTGGC**GATGATGTGTCTATTTCTCC |
| Ms_SepF_full_r | **GTTAGCAGCCGGATCT**CTACTTTCTAACTAAACTGACTCC |
| **Primers for colony PCR used during the construction of the pET-15b_*peiW* plasmid** | |
| T7 | TAATACGACTCACTATAGGG |
| T7_term | CTAGTTATTGCTCAGCGGT |
| **Primers for colony PCR used during the construction of the different pET-SUMO plasmids** | |
| 188_pAW-27 | CCCGCGAAATTAATACGACTCAC |
| 187_pAW-26 | CCTCAAGACCCGTTTAGAGGCC |
| *Overlaps for Gibson assembly are written in bold letters. Restriction sites are underlined. | |
